# Completing the TRB family: newly characterized members show ancient evolutionary origins and distinct localization, yet similar interactions

**DOI:** 10.1101/2022.11.23.517682

**Authors:** Alžbeta Kusová, Lenka Steinbachová, Tereza Přerovská, Lenka Záveská Drábková, Jan Paleček, Ahmed Khan, Gabriela Rigóová, Zuzana Gadiou, Claire Jourdain, Tino Stricker, Daniel Schubert, David Honys, Petra Procházková Schrumpfová

## Abstract

Telomere repeat binding proteins (TRBs) belong to a family of proteins possessing a Myb-like domain which binds to telomeric repeats. Three members of this family (TRB1, TRB2, TRB3) from *Arabidopsis thaliana* have already been described as associated with terminal telomeric repeats (telomeres) or short interstitial telomeric repeats in gene promoters (*telo*-boxes). They are also known to interact with several protein complexes: telomerase, Polycomb repressive complex 2 (PRC2) E(z) subunits and the PEAT complex (PWOs-EPCRs-ARIDs-TRBs). Here we characterize two novel members of the TRB family (TRB4 and TRB5). Our wide phylogenetic analyses have shown that TRB proteins evolved in the plant kingdom after the transition to a terrestrial habitat in Streptophyta, and consequently TRBs diversified in seed plants. TRB4-5 share common TRB motifs while differing in several others and seem to have an earlier phylogenetic origin than TRB1-3. Their common Myb-like domains bind long arrays of telomeric repeats in vitro, and we have determined the minimal recognition motif of all TRBs as one *telo*-box. Our data indicate that despite the distinct localization patterns of TRB1-3 and TRB4-5 in situ, all members of TRB family mutually interact and also bind to telomerase/PRC2/PEAT complexes. Additionally, we have detected novel interactions between TRB4-5 and EMF2 and VRN2, which are Su(z)12 subunits of PRC2.

## Introduction

Telomere repeat binding proteins (TRBs) were originally characterized as proteins in *Arabidopsis thaliana* with a binding affinity to telomeric DNA sequences proportional to the number of telomeric repeats (Schrumpfová et al. 2004). They belong to the plant-specific Single myb histone 1 (SMH) family with an N-terminal Myb-like domain (Myb-like), a central histone-like (H1/5-like) domain, and a coiled-coil domain near the C-terminus (Marian et al. 2003). Three members of the TRB family (TRB1, TRB2 and TRB3) in *A. thaliana* exhibit self-interactions and mutual interactions in the yeast-two hybrid (Y2H) system (Kuchař and Fajkus 2004; Schrumpfová et al. 2004). They bind plant telomeric repeats (TTTAGGG)_n_ through the Myb-like domain (Mozgová et al. 2008), while the H1/5-like domain is responsible for dimerization with other TRB proteins (Schrumpfová et al. 2008).

TRB1-3 proteins are proposed to participate in telomerase biogenesis. They interact directly with the catalytic protein subunit of telomerase (TERT) (Schrumpfová et al. 2014) and mediate interactions between TERT and Recombination UV B - like (RUVBL) proteins (Schořová et al. 2019), homologs of the essential mammalian telomerase assembly components Pontin and Reptin (Venteicher et al. 2008). Nuclear and predominantly nucleolar localization of TRB1-3 interacting with TERT and RUVBLs, as well as with the plant ortholog of dyskerin, CBF5 (Lermontova et al. 2007), was observed using Bimolecular fluorescence complementation (BiFC) (Sweetlove and Gutierrez 2019). Moreover, the TRB1 protein interacts via its H1/5-like domain with Protection of telomeres 1 (POT1b) (Schrumpfová et al. 2008), an *A. thaliana* homolog of the G-overhang binding protein Pot1, a core component of mammalian telomere cap complex, Shelterin (Tani and Murata 2005; Surovtseva et al. 2007). Additionally, in situ co-localization of TRB1 with telomeric DNA repeats has been detected in plant cells (Schrumpfová et al. 2014; Dreissig et al. 2017).

Telomere shortening was observed in *trb1* and *trb3* knockout mutants, in the *A. thaliana* ecotype Columbia with otherwise-stable telomere lengths (Shakirov and Shippen 2004; Schrumpfová et al. 2014; Zhou et al. 2016). In contrast, both telomere extension (Lee and Cho 2016) or shortening (Zhou et al. 2018) were detected in *trb2* knockout mutants of the *A. thaliana* ecotype Wassilewskija, which exhibits telomere length polymorphism in wild-type plants (Shakirov and Shippen 2004; Maillet et al. 2006). *Oryza sativa* with mutations in the *TRB1* gene also exhibited markedly longer telomeres, accompanied by genome instability (Hong et al. 2007).

In multicellular organisms, Polycomb repressive complex 1 (PRC1) and PRC2 repress target genes through histone modification and chromatin compaction. In *Drosophila melanogaster*,four core PRC2 subunits are present: the histone methyltransferase Enhancer of zeste [E(z)], Suppressor of zeste 12 [Su(z)12], Extra sex combs (Esc), and the histone-binding nucleosome remodelling factor 55 kDa (Nurf55). The E(z) homologs in *A. thaliana*, named CURLY LEAF (CLF) and SWINGER (SWN), are implicated in sporophyte development (reviewed in Mozgova and Hennig 2015). The PRC2 complex primarily methylates histone H3 on lysine 27 (H3K27me3), a mark of transcriptionally silent chromatin. TRB1–3 interact with the PRC2 proteins CLF and SWN (Zhou et al. 2018). We have shown that TRB1 proteins are not only associated with long arrays of telomeric repeats but also with interstitially located short telomeric sequences *telo*-box motifs, especially in the promoters of translation machinery genes (Schrumpfová et al. 2016). It was further shown that these *telo*-boxes are part of the cis-regulatory elements that may relate to PRC2 recruitment (Zhou et al. 2016; Zhou et al. 2018).

Besides the PRC2 complex, TRB1-3 are components of the PEAT complex (PWOs-EPCRs-ARIDs-TRBs) mediating histone acetylation/deacetylation and heterochromatin condensation. They potentially regulate the RNA-directed DNA methylation (RdDM) pathway (Tan et al. 2018; Tsuzuki and Wierzbicki 2018). The involvement of TRB proteins in histone deacetylation supports the previous observation that TRB2 directly interacts with histone deacetylases (HDT4 and HDA6) (Lee and Cho 2016). Recently, it has also been proposed that reciprocal binding of histone 1 (H1) and TRB1 to clustered *telo-box* motifs prevents H3K27me3 accumulation on large chromosomal blocks, e.g. telomeres (Teano et al. 2020).

Here we show that the plant-specific TRB proteins can be recognized in lower plants, such as Streptophytic algae, as well as in higher plants. In seed plants, TRB proteins are divided into three main lineages. We speculate that due to whole genome duplication (WGD) in *A. thaliana*, three ancestral TRB proteins have multiplied to the current five TRB members. We characterize new members of TRB family in *A. thaliana* (TRB4 and TRB5) and demonstrate that all members of the TRB family can bind long arrays of telomeric repeats with high specificity. We defined the minimal recognition motif for all TRBs as one *telo*-box. Even though TRB4 and TRB5 share very high sequence and structural homology with TRB1, TRB2 and TRB3, they differ in terms of the surface of their Myb-domains and their cellular localization. We provide evidence that TRB4-5 mutually interact with other members of the TRB family and physically interact with TERT, POT1a/b, SWN/CLF, and PWO1-3. Novel interactions were also detected between TRB4-5 and EMF2/VRN2, which represent the Su(z)12 subunits of PRC2. Completing TRB family analysis permits further exploration of the biological roles of these important plant-specific proteins.

## Results

### Sequence and Structural Divergences in the TRB family

Two decades ago, in silico analysis already predicted that the *A. thaliana* SMH family contains five TRB members (Marian et al. 2003). We used a combination of recent genome and transcriptome annotations (Lamesch et al. 2012; Cheng et al. 2017) and predicted sequence and structural similarity to characterize the TRB family members TRB4 and TRB5. These were each found to contain an N-terminal Myb-like domain, a central H1/5-like domain, and a C-terminal coiled-coil domain, similar to previously characterized family members TRB1-3 (Fig. 1A) (Marian et al. 2003; Schrumpfová et al. 2004; Mozgová et al. 2008).

**Fig. 1.**
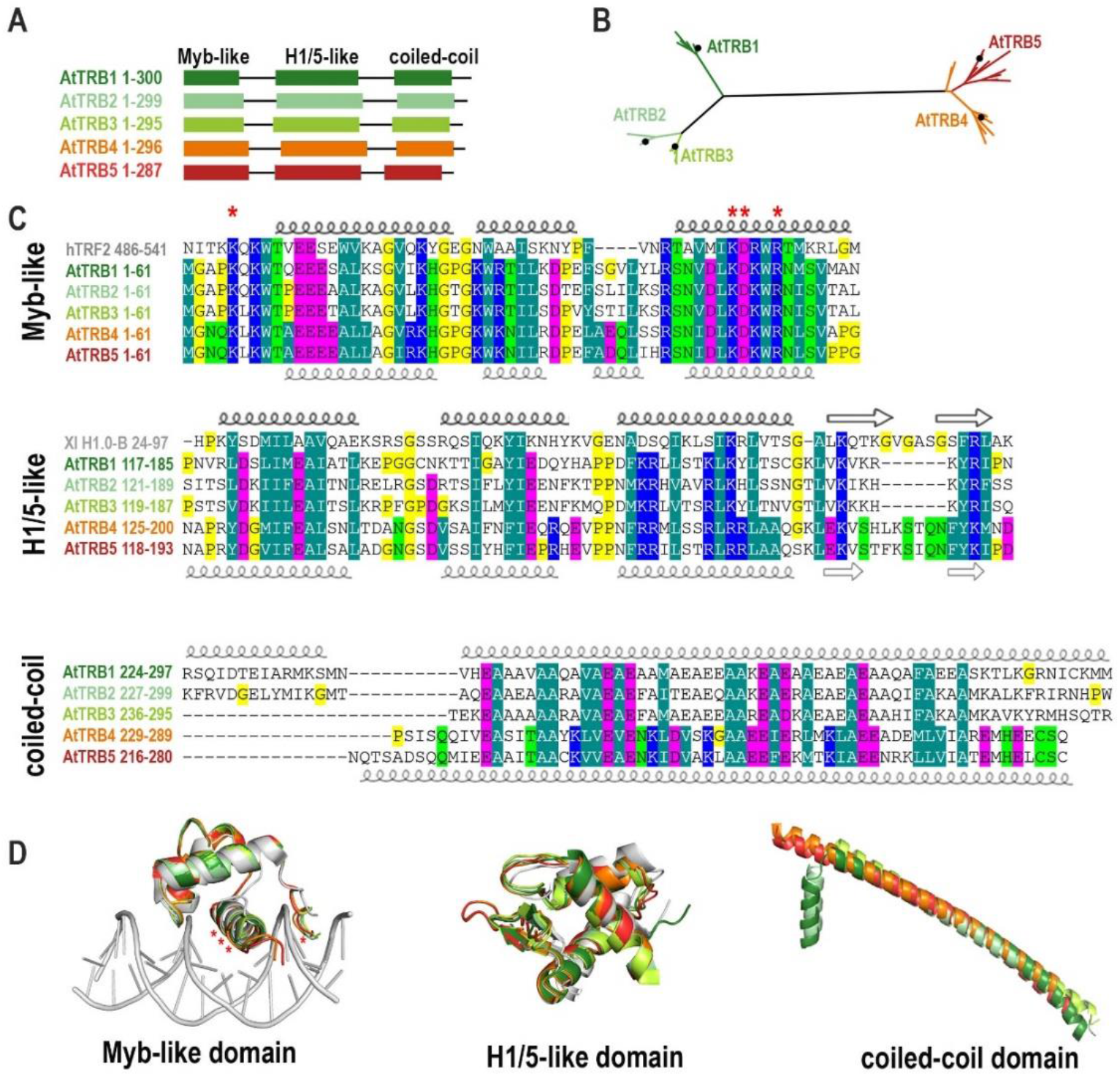
Sequence and structural alignments of TRB family proteins. A) Schematic representation of the conserved domains of TRBs from *A. thaliana. Myb-like*, Myb-like domain; *H1/5-like*, histone-like domain; *coiled-coil*, C-terminal domain. B) Unrooted Maximum likelihood (ML) phylogenetic tree of *Brassicaceae* TRB proteins. The length of the branches are proportional, and the black dots indicate the position of TRB1-5 from *A. thaliana*. For a list of species, see Supplementary Table 1. C) Multiple alignments of the Myb-like, H1/5-like and coiled-coil domains. The positions of α-helices or β-sheets of the uppermost or the lowermost sequence in each alignment are highlighted: *bold*, experimentally determined structures (cryo-EM or X-ray crystallography); *thin*, AlphaFold prediction. Human Telomeric repeat-binding factor 2 (hTRF2) and *Xenopus laevis* histone H1.0 (Xl H1.0-B) were used to show the most conserved amino acid (aa) residues. Amino acid shading indicates the following conserved amino acids: *dark green*, hydrophobic and aromatic; *light green*, polar; *blue*, basic; *magenta*, acidic; *yellow*, without side chain (glycine and proline). The aa of hTRF2 that mediate intermolecular contacts between telomeric DNA and hTRF2 are marked with an asterisk. D) Structural models of Myb-like, H1/5-like and coiled-coil domains. AlphaFold protein structure predictions deposited in the EMBL database were used (Varadi et al. 2022). The three-dimensional model of the Myb-like domain fits best the hTRF2-DNA interaction structure (PDB: 1WOU) (Court et al. 2005). The structure of the histone-like domain is most similar to *X. laevis* histone H1 structure (PDB: 5NL0) (Bednar et al. 2017). The positions of the aa of hTRF2 that mediate intermolecular contacts between telomeric DNA and hTRF2 are marked with an asterisk.

We then performed a phylogenetic reconstruction of TRB proteins from the *Brassicaceae* family based on a matrix including 52 TRBs, with 357 aligned positions. The tree shows that in *A. thaliana*, as well as in other *Brassicaceae*, TRB4 and TRB5 are grouped in a monophyletic lineage that is distant from TRB1-3 (Fig. 1B, Supplemental Table 1).

The Myb-like domain is composed of a helix-turn-helix (HTH) motif (Ogata et al. 1992; Bilaud et al. 1996). Even though the domain composition of SMH proteins is unique to the plant lineage, the Myb-like domain shows high aa sequence conservation across the plant and animal kingdoms (Fig. 1C). Predicted three-dimensional (3D) models of *A. thaliana* TRB Myb-like structural features were overlaid with the X-ray diffraction-resolved crystal structure of human Telomeric repeat-binding factor 2 (hTRF2) (Fig. 1D) and it was found that the 3D structure of the Myb-like domain is well conserved in both plant and animal kingdoms.

Interestingly, all TRBs show a slight difference in the composition of the second helix. However, the aa residues mediating the interaction between hTRF2 and the human telomeric repeat sequence (TTAGGG) are fully conserved in *A. thaliana* even though the plant telomeric sequence slightly differs (TTTAGGG) from the human one (Fig. 1C, D).

The central linker histone globular domain (H1/5-like domain) adopts a winged-helix fold including a HTH motif and a “wing” defined by two β-loops (Ramakrishnan et al. 1993). Using the SWISS-MODEL tool, the *Xenopus laevis* (Xl) H1 domain (XlH1.0) appeared to be the closest template to *A. thaliana* TRBs H1/5-like domain (Bednar et al. 2017). We compared the 3D structure of the histone globular domain from XlH1.0-B, obtained by cryo-electron microscopy (Cryo-EM) and X-ray crystallography (Bednar et al. 2017), with the AlphaFold protein structure predictions of the H1/5-like domain of *A. thaliana* TRBs (Varadi *et al.*, 2022). Although the sequence conservation between the XlH1 domain and the H1/5-like domain of *A. thaliana* TRBs is lower than in the Myb-like domain, the 3D structure of the H1/5-like domain in TRBs is highly similar (Fig. 1C, D).

The C-terminal coiled-coil domains usually contain a repeated pattern of hydrophobic and charged aa residues, referred to as a heptad repeat (Lupas and Gruber 2005). This repeating pattern enables two helices to wrap/coil around each other. The conservation of hydrophobic residues (Alanine (A), Isoleucine (I), Valine (V), Phenylalanine (F) or Methionine (M)) and an acidic residue (Glutamic acid (E)) is obvious in all coiled-coil domains from *A. thaliana* TRBs. Only one long α-helix is predicted by AlphaFold in TRB4-5, while the coiled-coil domains from TRB1 and TRB2 seem to have an additional short α-helix (Fig. 1C, D). However, these predictions await experimental verification.

The most divergent of the three domains in TRBs (Myb-like, H1/5-like and coiled-coil) is the coiled-coil domain. Our alignments suggest that although TRB4 and TRB5 are distant from TRB1-3 members in terms of sequence, they are folded into similar three-dimensional structures, with only minor differences.

### The Evolution of TRBs within the Plant Lineage

We performed comprehensive phylogenetic analysis to investigate whether TRBs are conserved across lower and higher plants, and whether TRB4-5 form a distinct group to TRB1-3 in all species, as observed in the higher plant *A. thaliana*. We used a data set of 268 proteins and 599 aligned positions and found that TRB proteins first evolved in Streptophyta in *Klebsormidiophyceae*, although TRBs are missing in the other Streptophyte algae studied, including *Charophyceae* and *Zygnematophyceae*. In *Klebsormidium nites* only one TRB homolog was identified. Following the evolutionary tree, an increasing number of TRB homologues were found in Bryophyta and Tracheophyta. There are three homologues of TRB in mosses *Sphagnum fallax* and *Physcomitrium patens*, and these TRBs share very high sequence similarities. Contrastingly, there is only one homologue in the moss *Ceratodon purpureus*. Perhaps due to limited sequence information available for the genomes of hornworts or liverworts, TRB was not found in these lineages. In seed plants, which have undergone more rounds of whole genome duplication events (WGDs) than Bryophyta and Lycophyta (Clark and Donoghue 2018), predominantly three TRB proteins were recognized. Within *Brassicaceae*, which has undergone an additional recent round of WGD (Walden et al. 2020), five TRB homologs were revealed (Fig. 2A).

**Fig. 2.**
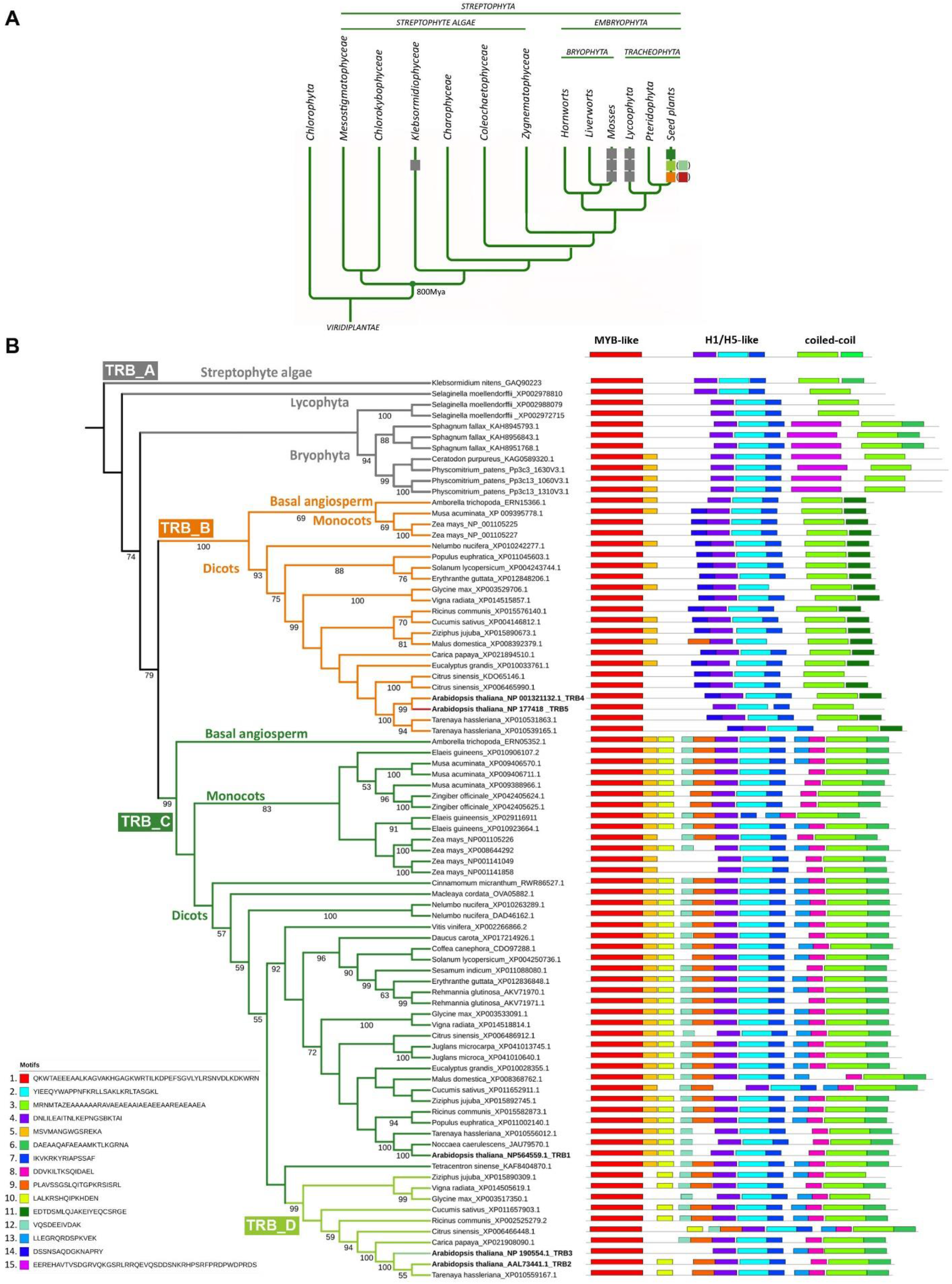
Phylogenetic analysis of TRB proteins. A) Simplified evolution of the main Viridiplantae lineages with known TRB proteins. One TRB protein evolved initially in Streptophyta in *Klebsormidiophyceae*, then diversified to three similar homologues in Mosses and Lycophyta and several diverse homologues in seed plants, with five members in *A. thaliana* from *Brassicaceae*. The evolutionary tree was adopted from Rensing 2020 and Cheng et al. 2019. B) ML phylogenetic tree of TRB proteins. Twenty-seven species were included in 83 sequences with 465 bp in the final data set. Only one species from the family was selected for the final analysis. The ML likelihood is −24356.573420. The numbers below branches indicate bootstrap support values >50%. Four major groups are shown in the phylogenetic tree: TRB_A for Streptophyte algae, Lycophyta and Bryophyta; TRB_B including AtTRB4 and AtTRB5 for Embryophyta; TRB_C for Embryophyta with AtTRB1 and the nested TRB_D encompassing AtTRB2 and AtTRB3. Motifs are ranked and ordered by the highest probability of occurrence. Fifteen most probable motifs are depicted. For sequence and sequence conservation information, see Supplementary Fig. 1. For a list of species, see Supplementary Table 2.

Next, the MEME search (Bailey et al. 2009) was used to identify the 15 most typical motifs in TRBs across the plant kingdom (Supplemental Fig. 1). For ease of presentation, a simplified tree was used with 83 representatives from 33 families and 26 orders, each family having only one member (Fig. 2B, Supplemental Table 2). TRB proteins were divided into three main lineages named TRB_A, TRB_B and TRB_C, into the latter of which the TRB_D sub-lineage is integrated. The lineage TRB_A includes Streptophyte algae, Bryophyta and Lycophyta and diverges from seed plant lineages TRB_B and TRB_C in terms of the length of introns, disordered parts of proteins or additional motifs. A clear diversification of TRBs into monocots and dicots was revealed in both TRB_B and TRB_C lineages. In seed plants, the TRB_B lineage seems to be less abundant than in the TRB_C lineage.

Consistent with previous findings, the canonical N-terminal Myb-like (motif 1), a central H1/5-like (motifs 2, 4 and 7), and a C-terminal coiled-coil (motif 3) domains were amongst the top conserved motifs present in all TRBs (Fig. 2B). Within the lineage TRB_A, a unique aa motif was detected in Bryophytes (motif 15; EEREH). Interestingly, the coiled-coil domains of the proteins from TRB_B lineage contain the specific motif 11 (EDTDS), but most of the proteins from the TRB_A and TRB_C lineage contain the motif 6 (DAEAA) instead. However, motif 6 is not present in the *A. thaliana* protein TRB5 that has diverged from TRB4. The *A. thaliana* proteins TRB4 and TRB5 (belonging to the TRB_B lineage) lack motif 8 (DDVKI) adjacent to the coiled-coil domain. In contrast, this motif is present within proteins of the TRB_C lineage, including *A. thaliana* TRB1-3 proteins.

The TRB_D sub-lineage is embedded within TRB_C lineage but supported by a high 99% bootstrap value. TRB2 and TRB3 from *A. thaliana* are nested within the TRB_D sub-lineage. This lineage lacks motif 5 (MSVMA), a motif adjacent to the Myb-like domain, which is present in the rest of the TRBs in the TRB_C lineage. Similarly to *Brassicaceae*, divergence to the TRB_D sub-lineage was detected within several other dicot families (e.g., *Rhamnaceae - Ziziphus jujuba, Cucurbitaceae - Cucumis sativus, Euphorbiaceae - Ricinus communis, Rutaceae - Citrus sinensis, Malvaceae - Glycine max, Cleomaceae* - *Tarenaya hassleriana*).

In general, TRBs were detected in lower and higher plants. TRBs in Streptophyte algae, Lycophyta and Bryophyta (grouped in TRB_A lineage) are more closely related to *A. thaliana* TRB4 and TRB5 proteins (TRB_B lineage) than to *A. thaliana* TRB1_3 (TRB_C lineage). Lineage TRB_B and TRB_C differ in several motifs accompanying the canonical H1/5-like or coiled-coil domains. The specificity of the sub-lineage TRB_D, embedded into TRB_C lineage, is highlighted by its lack of a specific motif 5 following the canonical Myb-like domain.

### Dicots from TRB_B lineage differ solution accessible surface of the Myb-like domain

The N-terminal Myb-like domain is the most conspicuous structural unit in TRBs. In order to compare the structural characteristics of this, we further examined the sequence conservation and estimated evolutionary conservation of predicted 3D structures. The TRB groups (TRB_A, B, C, D), established in Fig. 2B, were subdivided into Lycophytes, Bryophytes, Monocots and Dicots. Consensus sequences of the Myb-like domain in each individual group are visualized in Fig. 3A.

**Fig. 3.**
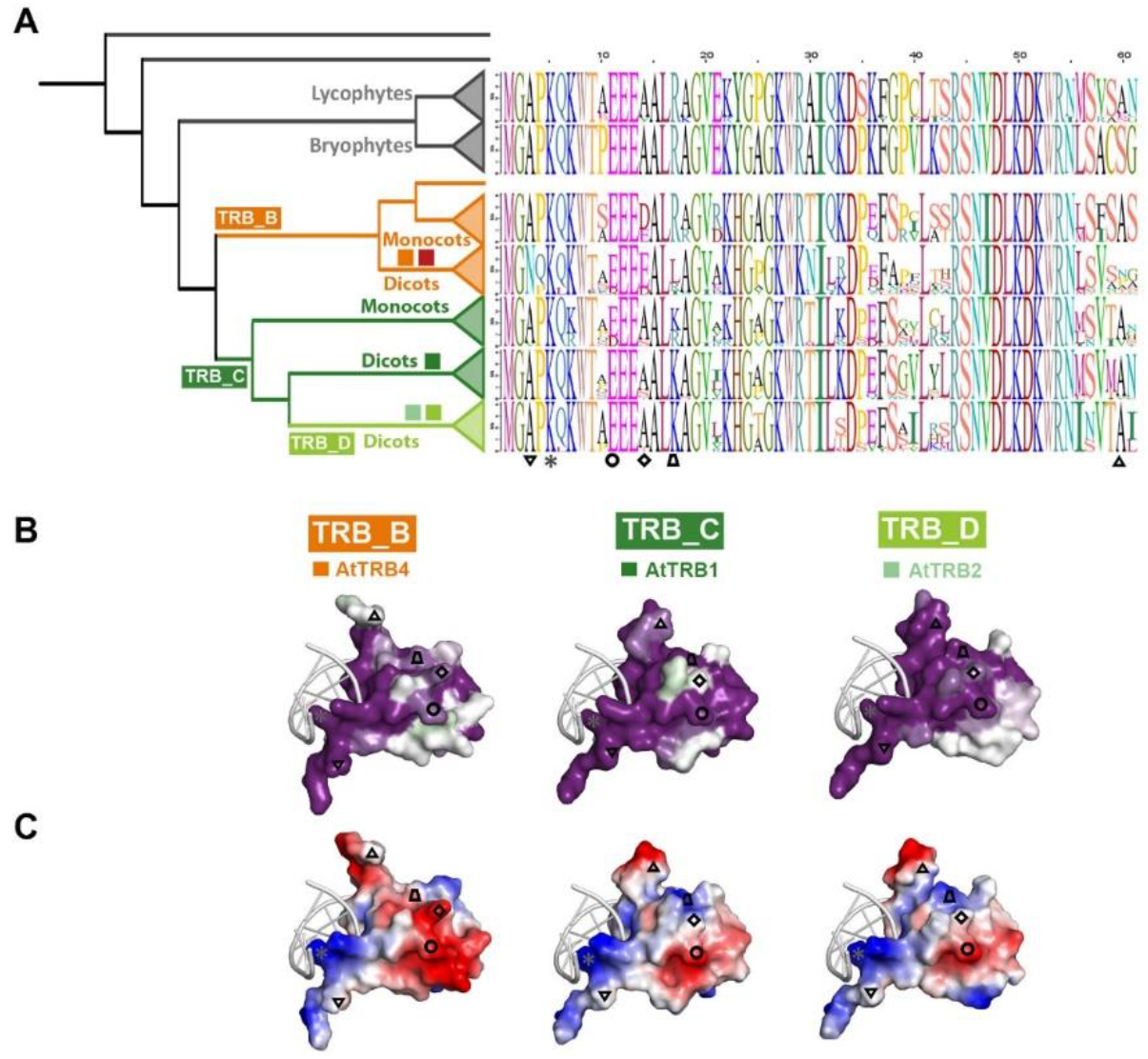
Conserved features of MYB-like domain through TRB lineages. A) Consensus sequences of Myb-like domain in each evolutionary lineage were visualised using sequence logos. *Colored squares*, represent TRB1-5 proteins from *A. thaliana; Inverted triangle*, the aa at position 3 (A) at N-terminus of TRBs that is replaced in Dicots from TRB_B lineage to N; *Asterisk*, the aa at position 5 (K) that mediates intermolecular contacts between telomeric DNA and N-terminus of all TRBs (Fig. 1C, D); *Circle*, the negatively charged aa (E/D) at position 11 conserved across all TRB lineages; *Rhombus*, the aa at position 14 replaced from uncharged (A/S) to negatively charged (E/D) in the whole TRB_B lineage; *Trapezoid*, the aa at position 17 that is in Dicots from TRB_B lineage partially replaced from positively charged (K/R) to hydrophobic (L); *Triangle*, the aa at position 60 that shows lower conservation in Dicots from TRB_B lineage than in other lineages. B) The evolutionary dynamics of aa substitutions among aa residues within Dicots in TRB_B, TRB_C and TRB_D lineages, exemplified in *A. thaliana* TRB4, TRB1 and TRB2, respectively, were visualized using ConSurf 2016 (Ashkenazy et al. 2016). The conservation of residues is presented in a scale, where the most conserved residues are shown in dark magenta and non-conserved residues as white. C) Surface models showing the charge on the Myb-like domain are presented from the site opposing the DNA-binding viewpoint for each model. Residue charges are coded as red for negative, blue for positive, and white for neutral, visualised using PyMol, Version 2.4.1, Schrödinger, LLC.

The evolutionary dynamics of aa substitutions in *A. thaliana* TRB4, 1 and 2, representing Dicots from lineages TRB_B, C and D, respectively, were visualized using ConSurf 2016 (Ashkenazy et al. 2016). Visualization has shown a very strong conservation of the DNA-binding surface in all lineages (Supplemental Fig. 2), consistent with the above analysis (Figs. 1 and 2). In comparison, opposite solution-accessible surface was less conserved; in Dicots from the TRB_D lineage, conservation was lower than in Dicots from TRB_B and TRB_C lineages (Fig. 3B, Supplemental Fig. 2). Dicots in all three lineages exhibit a variant unstructured N-terminus of the MYB-domain which is conserved within the lineage. The aa at the N-terminal position 3 in Dicots from the TRB_B lineage, including TRB4-5 from *Arabidopsis*, show significantly conserved substitutions of Alanine (A) residues to the polar uncharged aa Asparagine (N) (Fig. 3, *Inverted triangle*). The unstructured C-terminal tail of the Myb-like domain in the TRB_B lineage (Fig. 3, *Triangle*, Supplemental Fig. 2A) shows less conservation than in TRB_C and TRB_D lineages, where these residues are part of the third helix.

Surface models showing the charge of the Myb-like domain were visualized using PyMol viewer (Fig. 3C, Supplemental Fig. 2C, F). In the TRB_B lineage, visualizations reveal a negatively charged aa (E/D) at position 14 (Fig. 3, *Rhombus*) flanking conserved EEE motif (Fig. 3, *Circle*), whereas proteins in the TRB_A and TRB_C lineages possess uncharged Alanine (A) at the position 14. Interestingly, the aa (positively charged aa K/R at position 17) proximal to the E/D motif, are replaced to uncharged (L) in Dicots from TRB_B lineage (Fig. 3C, *Trapezoid*).

These data indicate that the E/D motif at position 14 together with aa at position 17 are responsible for the additional areas of negative charge on the solution-accessible surface of the Myb-domain in Dicots from TRB_B lineage.

### Even one telomeric unit is sufficient for TRB binding

Our previous findings revealed that the N-terminal Myb-like domains of TRB1-3 are responsible for specific recognition of long arrays of telomeric DNA (Schrumpfová et al. 2004; Mozgová et al. 2008). The DNA-binding preference for oligodeoxynucleotide (oligo) substrates was tested using the electrophoretic mobility shift assay (EMSA). Oligo sequences were designed to assess the effect of TRB4 and TRB5 binding to long arrays of telomeric sequences as well as to interstitially located *telo*-boxes (Fig. 4A, Supplemental Table 4). Either a tetramer of telomeric sequences or a *telo*-box sequence (1.2 telomeric units) flanked with non-telomeric DNA were used as the labelled probes. A non-telomeric oligonucleotide, added in 1, 20 and 100-fold excess, served as competitor DNA and *vice versa*. Parallel experiments with the TRB1 protein were used for comparison.

**Fig. 4.**
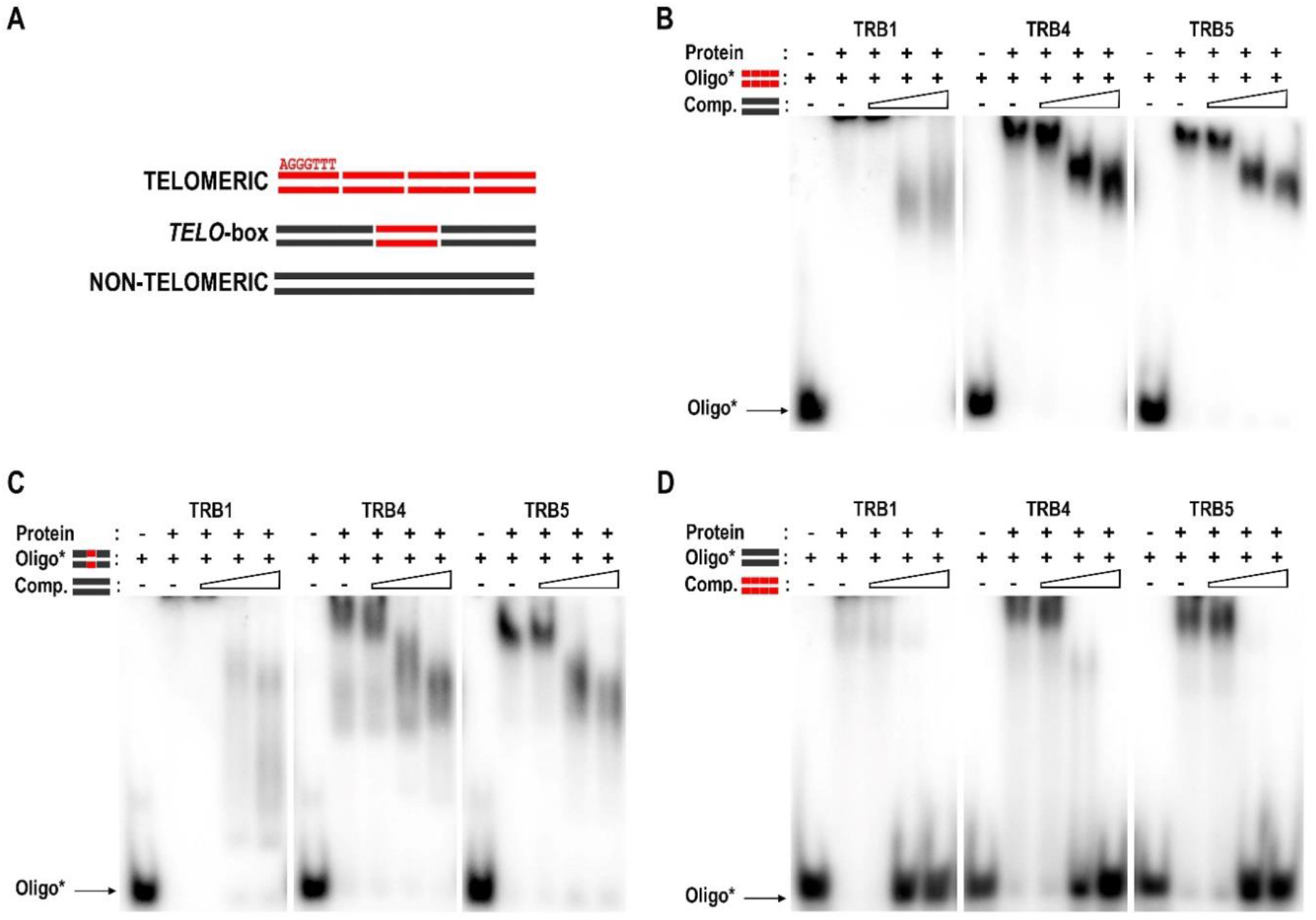
EMSA of TRB1, TRB4 and TRB5 binding of radioactively-labelled oligonucleotides. A) Schematic depiction of oligonucleotides employed. *Telomeric*, four repeats of plant telomeric DNA sequence; *telo-box*, 1.2 plant telomeric units flanked with non-telomeric DNA sequence; *non-telomeric*, oligonucleotide with non-telomeric DNA. B) EMSA of TRB1, TRB4 and TRB5 proteins binding radioactively-labelled double-strand (ds) tetramers of telomeric sequence with unlabelled tetramers of non-telomeric oligonucleotides as competitor DNA. The concentration of unlabelled competitor increases from 1-, 20- to 100-fold the concentration of the labelled probe (as depicted by the triangle). Oligo*: protein ratio is 1:10. C) EMSA of the same proteins with a radioactively-labelled ds *telo*-box oligonucleotides with unlabelled non-telomeric oligonucleotides as competitor DNA, performed as in B. D) EMSA of the same proteins with a radioactively-labelled ds of non-telomeric oligonucleotides with unlabelled ds tetramers of telomeric sequence as competitor DNA, performed as in B.

The results obtained from these experiments clearly show that TRB4 and TRB5 do indeed preferentially bind long arrays of telomeric sequences or *telo*-boxes positioned within a non-telomeric DNA sequence (Fig. 4B-D). TRBs bind to telomeric dsDNA in a similar mode as was described for TRB1-3, forming a high-molecular-weight complex which does not migrate into the gel (Schrumpfová et al. 2004; Mozgová et al. 2008). This complex is only partially disassembled by the addition of a 20 or 100-fold excess of the non-telomeric competitor, but the competitor did not release the protein from the complex with telomeric DNA. In contrast, the high-molecular-weight TRBs complexes with non-telomeric oligos were completely dissassambled upon the addition of a 20-fold excess of the telomeric competitor.

Our data indicate that TRB4-5, as well as other TRBs, are capable of binding long arrays of telomeric sequences or short motifs with as little as one telomeric repeat. This is in contrast to human TRF1/TRF2, which bind two telomeric repeats as preformed dimers (Court et al. 2005) and also to previous predictions (Hofr et al. 2009). Having defined the TRB minimal recognition motif as one *telo*-box, the binding properties of TRBs are poised for further investigation.

### Unlike other TRB family members, TRB5 is preferentially localized in the cytoplasm

To compare *Arabidopsis* TRB proteins further, we examined the subcellular localization of native TRB proteins in *Arabidopsis* cells, using a mouse monoclonal antibody developed in our laboratory specific for the conserved section of the Myb-like domain found in the TRB family. The anti-TRB 5.2 antibody recognizes all five members of TRB family (Schrumpfová et al. 2014). Nuclei isolated from 10-day-old seedlings were subjected to immunofluorescence using this anti-TRB antibody combined with DAPI staining. We observed a speckled distribution of TRBs in the nucleus and nucleolus (Fig. 5A).

**Fig. 5.**
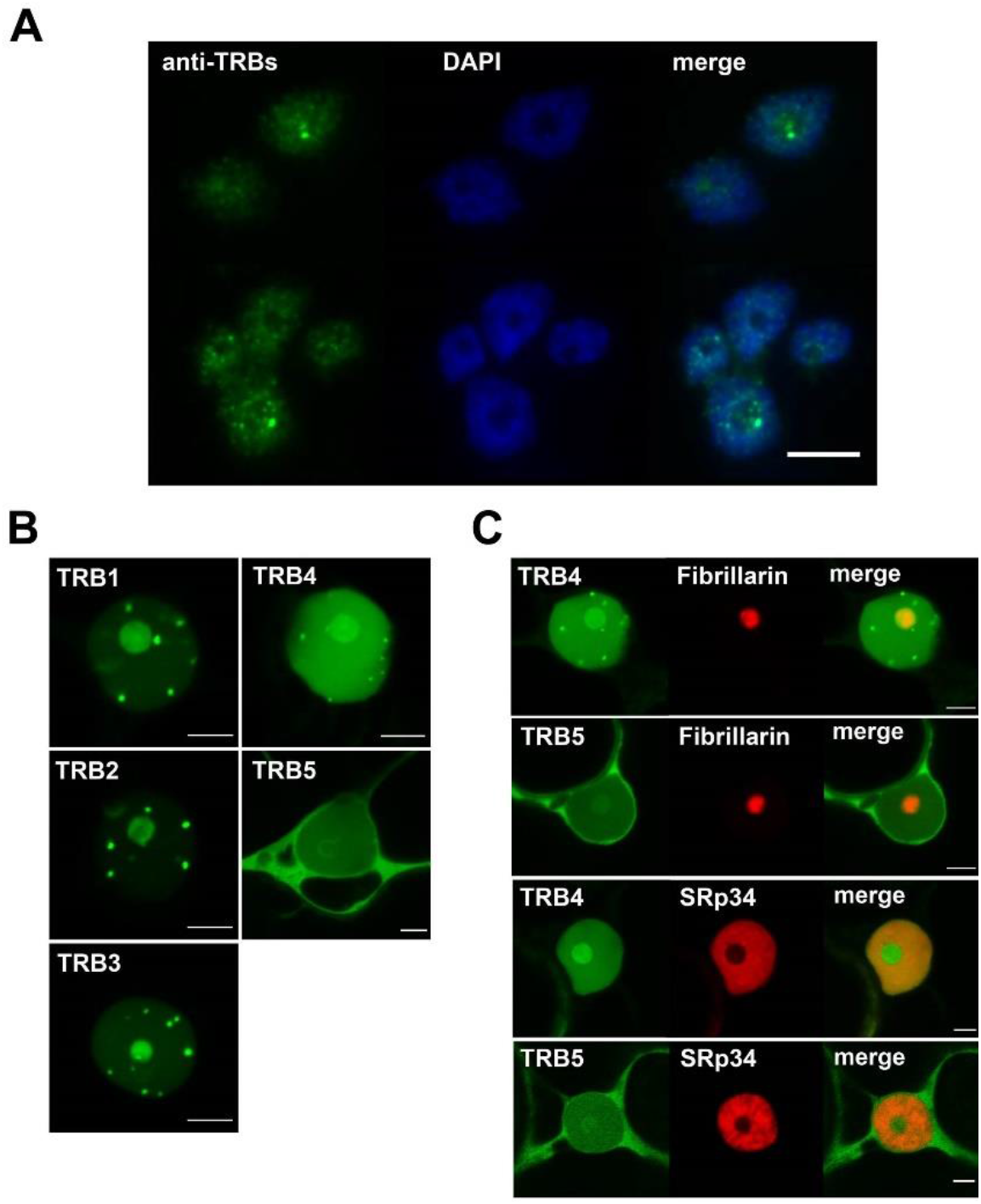
Subcellular localization of native TRBs and GFP-TRB fusion proteins. A) Isolated nuclei from *A. thaliana* seedlings were subjected to immunofluorescence using an anti-TRB antibody combined with DAPI staining. All five native members of the TRB family are visualized. Scale bar = 10 μm. B) TRB1-5 were fused with GFP (N-terminal fusions), expressed in *N. benthamiana* leaf epidermal cells and observed by confocal microscopy. Single images of areas with nuclei are presented. Scale bars = 5 μm. C) Co-localization of TRB4 and TRB5 (N-terminal GFP fusions) with a nucleolar marker (Fibrillarin-mRFP) and a nucleoplasm marker (SRp34-mRFP) was performed as described in B). Single images of areas with nuclei are presented. Scale bars = 5 μm.

As the anti-TRB 5.2 antibody does not distinguish the localization of individual proteins, we proceeded to the subcellular localization of individual members of TRB family using TRBs fused with green fluorescent protein (GFP). Confocal microscopy showed that TRB1-3 fused with GFP expressed from pGWB6 after transient *Agrobacterium*-mediated transformation in *N. benthamiana* leaf epidermal cells are localized mainly in nucleoli and nucleoplasmic fluorescence foci, as was described previously (Schrumpfová et al. 2014; Zhou et al. 2016). Interestingly, TRB4 is distributed not only in nucleoli and nucleoplasmic fluorescence foci of different sizes, but also throughout the nucleoplasm. Conversely, TRB5 fused with GFP is localized mainly in the cytoplasm with only minor localization in the nucleolus and nucleoplasm (Fig. 5B, Supplemental Fig. 3 and 4).

Co-expression of GFP-TRBs with the nucleolar marker Fibrillarin1 or nucleoplasm marker Serine-arginine-rich proteins 34 (SRp34) fused with a monomeric red fluorescent protein (mRFP) after transient *Agrobacterium*-mediated transformation in *N. benthamiana* leaf epidermal cells verified the subnuclear localizations of the TRB4 and TRB5 described above. Using Fibrillarin-mRFP we clearly identified the stronger localization of GFP-TRB4 in the nucleolus, but the weak nucleolar localization of GFP-TRB5. The relative positioning of GFP-TRB and the nucleoplasm marker SRp34-mRFP also supports our observation that the GFP-TRB5 is preferentially localized in the cytoplasm (Fig. 5C).

TRB4-5 fused with GFP show a distinct localization in the plant cell compared to TRB1-3, proteins with a later evolutionary origin. In particular, GFP-TRB5 manifests strong cytoplasmic localization, suggesting a possible specific functional role or spatiotemporal regulation.

### Dimerization of TRB proteins

To shed light on the conservation of mutual interactions between TRB proteins from *Arabidopsis*, we analyzed interactions between all members of TRB family. Do date, self and mutual dimerization of TRB1-3 has been investigated by Y2H or by co-immunoprecipitation experiments (Co-IP) (Kuchař and Fajkus 2004; Schrumpfová et al. 2004; Schrumpfová et al. 2008; Mozgová et al. 2008), however knowledge of the precise subcellular localization of mutual TRB interactions is missing.

First, the interactions of TRB4 and TRB5 with other TRB family members were investigated using a GAL4 based Y2H assay. Mutual interactions between TRB family members from *Arabidopsis* appear to be conserved, as both TRB4 and TRB5, interact with all TRB family members. Moreover, TRB4-5 form self-dimers in a similar way to TRB1-3 (Fig. 6A).

**Fig. 6.**
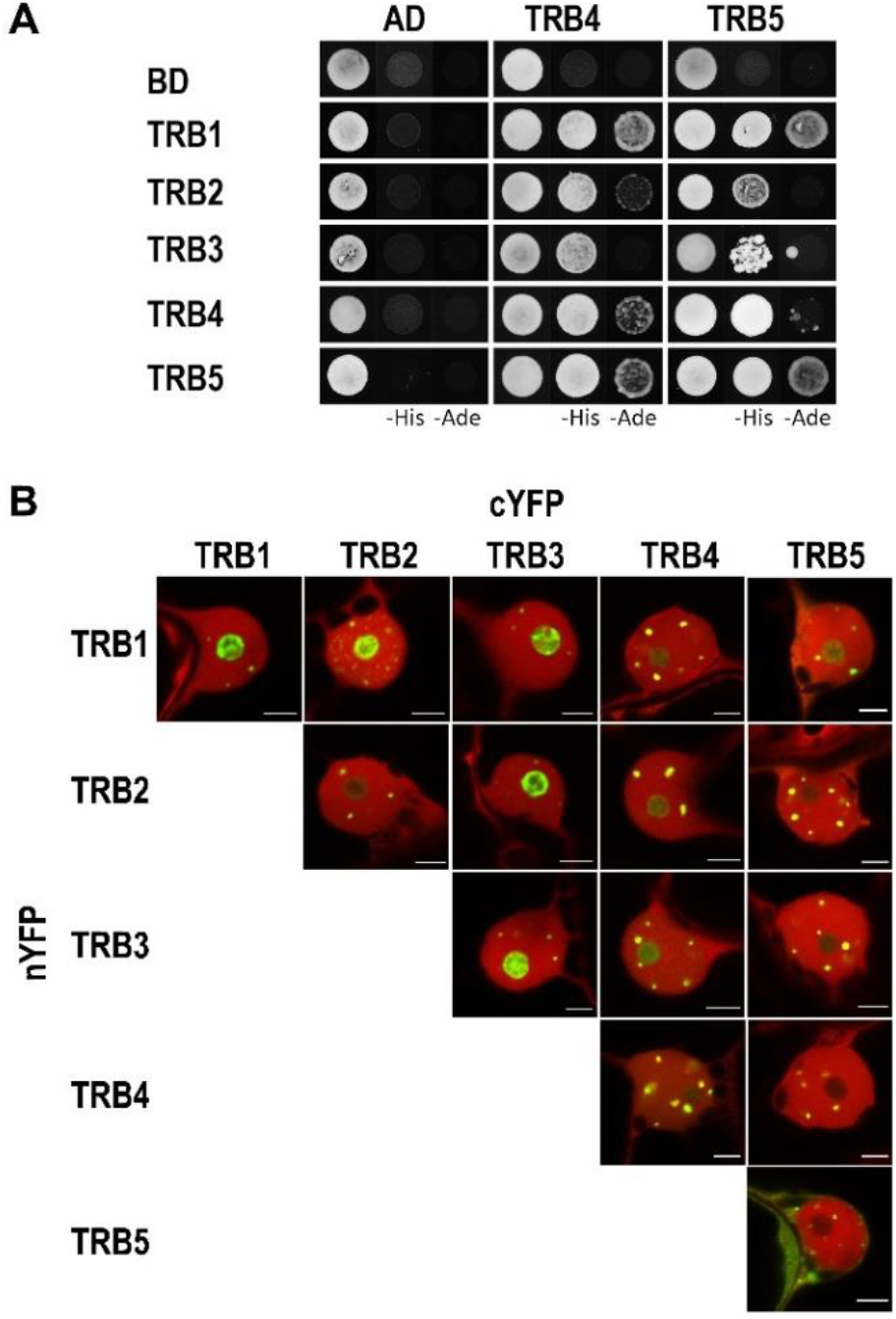
Dimerization of TRB proteins. A) The Y2H system was used to assess mutual proteinprotein interactions of TRBs. Two sets of plasmids carrying the indicated protein fused to either the GAL4 DNA-binding domain (BD) or the GAL4 activation domain (AD) were constructed and introduced into yeast strain PJ69-4a carrying reporter genes His3 and Ade2. Interactions were detected on histidine-deficient SD medium (-His), or under stringent adenine-deficient SD medium (-Ade) selection. Co-transformation with an empty vector (AD, BD) served as a negative control. B) Interactions of TRBs fused with nYFP or cYFP part were detected using the Bimolecular fluorescence complementation (BiFC) assay in *N. benthamiana* leaf epidermal cells. Shown here are single images of merged signals of reconstructed YFP (interaction of the tested proteins) and signals of mRFP (internal marker for transformation and expression) fluorescence detected by confocal microscopy. For separated fluorescent emissions, see Supplemental Fig. 5. Scale bars =5 μm.

Next, BiFC assays with equal protein levels were used to detect self- and mutual interactions of TRBs at the level of cellular compartments. The TRB coding sequences were introduced into 2in1 multiple expression cassettes within a single vector backbone with an internal marker for transformation and expression - mRFP1 (Grefen and Blatt 2012). TRBs were fused with an N- and/or C-terminal half of yellow fluorescent protein (nYFP, cYFP) as both N-terminal and C-terminal fusions. A list of all constructs is provided in Supplemental Table 3. Fluorescence was detected using confocal laser scanning microscopy after transient *Agrobacterium*-mediated transformation in *N. benthamiana* epidermal cells. Our observations show self- and mutual interactions of TRB1-4 in the nucleolus and/or in nucleoplasmic fluorescence foci that were of different numbers and sizes. Interestingly, the TRB5 homodimeric interaction was found to be clearly cytoplasmic with a reduced nuclear speckle size compared to other TRB interactions. Additionally, TRB1-3, but not TRB4, interact moderately with TRB5 in the nucleolus, as well as in more prominent nucleoplasmic fluorescence foci (Fig. 6B, Supplemental Fig. 5 and 6).

Overall, our characterization of mutual TRB interactions revealed that all TRB members have the ability to form self-dimers and mutually interact. However, TRB5 homodimeric interactions are predominantly localized in the cytoplasm, which differs from TRB1-4 homodimeric interactions, which are nucleolar or localized in nucleoplasmic fluorescence foci.

### Novel interaction partners of TRB proteins - PRC2 subunits EMF2 and VRN2

To further elucidate the conservation of protein interaction partners throughout the TRB family, we looked for interactions between TRB4 and TRB5 with TERT, RUVBLs, POT1b, PWO1-3, SWN, CLF interactors, which have already been described as interacting with TRB1-3 (Schrumpfová et al. 2008; Schrumpfová et al. 2014; Hohenstatt et al. 2018; Zhou et al. 2018; Tan et al. 2018; Schořová et al. 2019; Mikulski et al. 2019). These interactions were investigated using a GAL4-based Y2H assay, and TRB interactions with newly identified protein interactors were verified by Co-IP.

Our results revealed that TRB4-5 interact with the N-terminal domains of TERT called telomerase essential N-terminal (TEN, 1 – 233 aa) and RNA interaction domain 1 (RID1, 1 – 271 aa) fragments, in a similar manner to that described for TRB1-3 (Schrumpfová et al. 2014). Interactions of TRBs with TERT seem to be mediated only via the N-terminal domain of TERT, as the centrally positioned Telomerase RNA-binding domain (TRBD) or reverse transcriptase domain (RT) domain do not show interactions (Supplemental Fig. 7). We also observed interactions of TRB4-5 with RUVBL1, RUVBL2A and POT1b in the same manner as was observed for TRB1-3. Moreover, Y2H experiments detected novel interactions between TRB4-5 with a second homolog of the human POT1 protein in *A. thaliana* - POT1a. These interactions were further confirmed by the Co-IP of these proteins (Fig. 7A, Supplemental Fig. 8A).

**Fig. 7.**
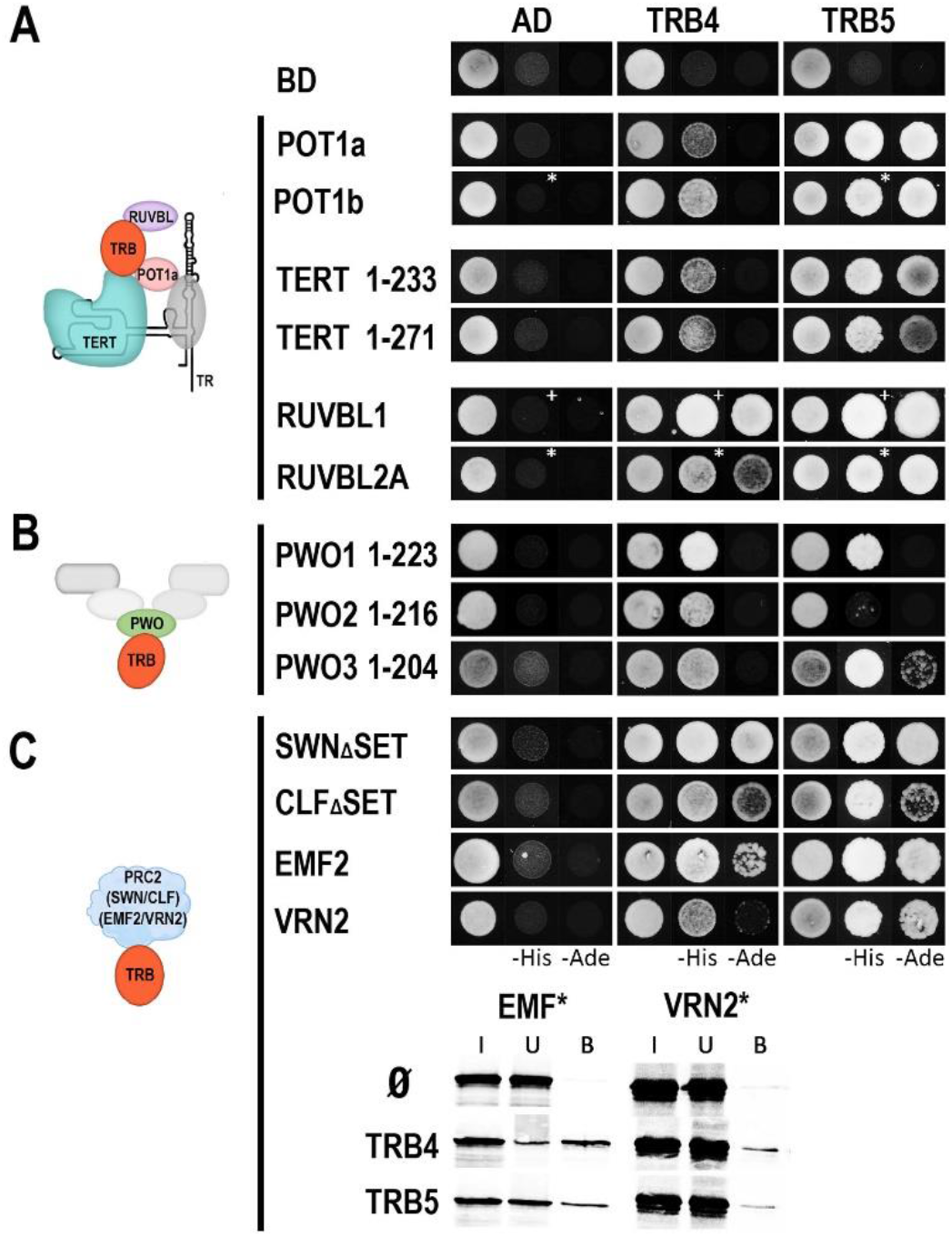
Interaction of TRB4-5 with various partners. A) The Y2H system was used to assess proteinprotein interactions of TRB4-5 proteins with TERT fragments, RUVBLs and POT1a/b as in Fig.6. Co-transformation with an empty vector (AD, BD) served as a negative control. *Asterisks*, 1 mM 3-aminotriazol (3-AT); *cross*, 3 mM 3-AT. B) Interactions between N-terminal domain of PWO1-3 and TRB4-5 were detected as in A). Interactions with full length PWO1-2 proteins are in Supplementary Fig. 8B. C) Novel interactions between TRB4-5 and EMF2/VRN2, as well as interactions with SWN_Δ_SET/CLF_Δ_SET were tested using Y2H system as in A). Novel interactions were verified by Co-IP. The TNT expressed VRN2 and EMF2 (^35^S-labelled*) were mixed with TRB4-5 (myc-tag) and incubated with anti-myc antibody. In the control experiment, the VRN2 and EMF2 proteins were incubated with anti-myc antibody and Protein G magnetic particles in the absence of partner protein. Input (I), unbound (U), and bound (B) fractions were collected and separated in SDS– 10% PAGE gels.

Furthermore, TRB4-5 interact with PWOs, members of the PEAT complex, as was already observed for TRB1-3 (Tan et al. 2018). A strong interaction was detected between TRB4-5 and PWO1 full-length (Supplemental Fig. 8B) or PWO1 N-terminal sections, including PROLINE-TRYPTOPHANE-TRYPTOPHANE-PROLINE (PWWP) domain (1-223 aa) (Fig. 7B). PWO2 and 3 interact with TRB4 via their N-terminal parts (1-216 and 1-204 aa, respectively). Similarly, we detected the interactions of the N-terminal parts of PWO1 and PWO3 with TRB5.

To identify TRB interactions with E(z) homologs, members of the PRC2 complex, we used SWN and CLF proteins without the SET [Su(var)3-9, E(z), Trx] domain that confers histone methyltransferase activity (SWN_Δ_SET and CLF_Δ_SET) (Chanvivattana et al. 2004; Hohenstatt et al. 2018). The SET-domain does not seem to be involved in interactions of SWN and CLF with TRB4-5 as Y2H experiments identified interactions between TRB4-5 and SWN_Δ_SET and CLF_Δ_SET (Fig. 7C).

It was shown recently that the rice homologue of TRB, Telomere repeat-binding factor 2 (TRBF2), and rice Su(z)12 homologues of the PRC2 complex named EMBRYONIC FLOWER 2b (EMF2b) interact with each other (Xuan et al. 2022). We observed novel interactions, not only between *Arabidopsis* TRB4-5 and EMF2, but also between TRB4-5 and other Su(z)12 homologue VERNALIZATION 2 (VRN2). The observed interactions from Y2H system were verified by Co-IP (Fig. 7C).

This broad screen of TRB4-5 protein interaction partners suggests a role of TRBs in various protein complexes. These complexes may be responsible for the recruitment of TRBs to telomeric DNA repeats (e.g., Telomerase complex to telomeres) (Fig. 8A) or to *telo*-boxes localized in the promoter regions of the various genes (e.g., PEAT complex or PRC2 subunits to *telo*-boxes) (Fig. 8C, D, F).

**Fig. 8.**
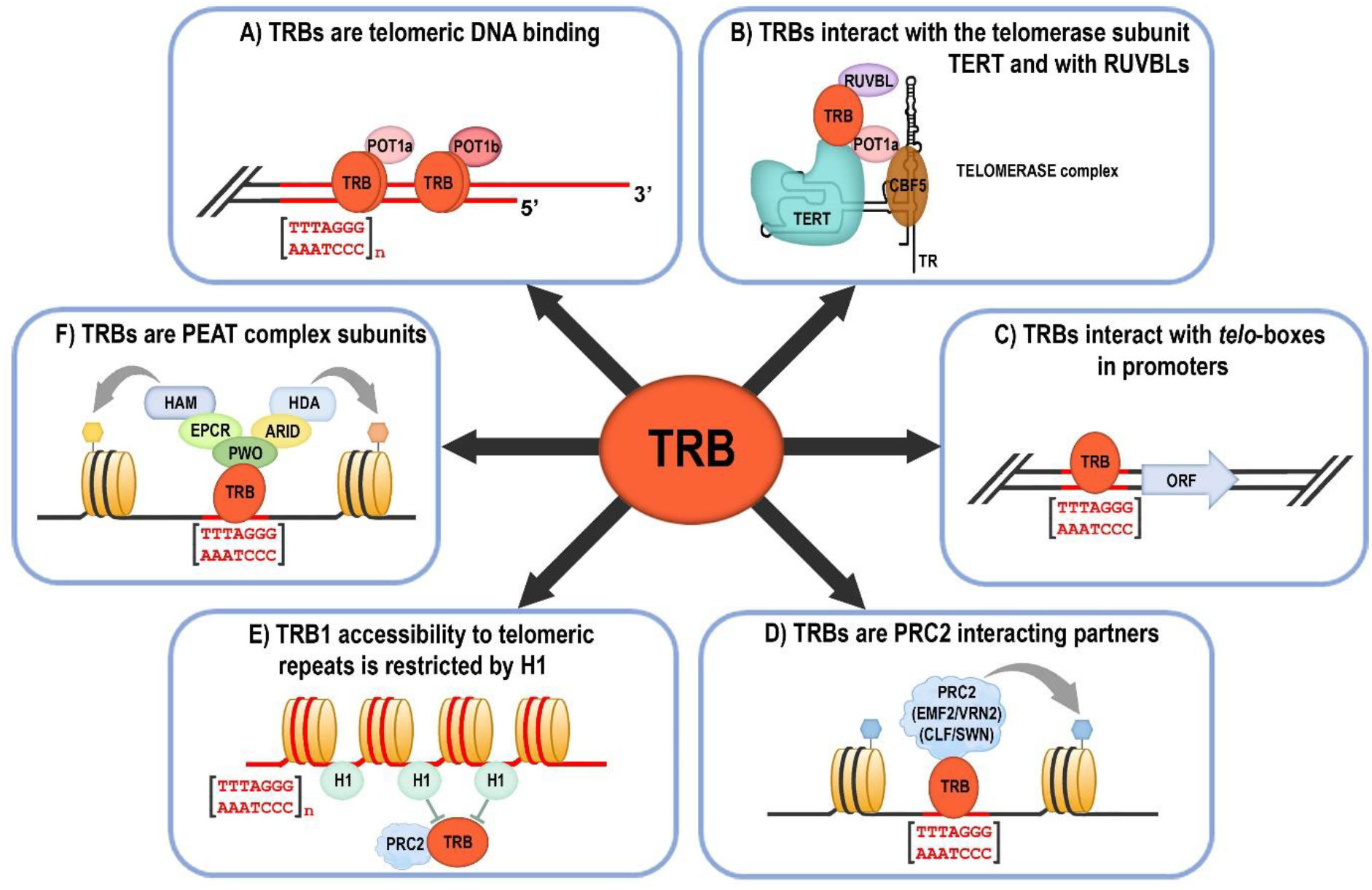
Overview of the main Telomere repeat binding proteins (TRBs) functions. A) TRBs are associated with the physical ends of chromosomes (telomeres) via their Myb-like domain (Schrumpfová et al. 2004; Mozgová et al. 2008; Dvořáčková et al. 2010; Schrumpfová et al. 2014; Dreissig et al. 2017). TRBs interact with Arabidopsis homologs of the G-overhang binding protein Protection of telomere 1a, b (POT1a, b) (Schrumpfová et al. 2008, this study). B) TRBs mediate interactions of Recombination UV B – like (RUVBL) proteins with the catalytic subunit of telomerase (TERT), and participate in telomerase biogenesis (Schrumpfová et al. 2014; Schořová et al. 2019). TRBs are associated in the nucleus/nucleolus with POT1a (Schořová et al. 2019), and also with a plant orthologue of dyskerin, named CBF5 (Lermontova et al. 2007) that binds the RNA subunit of telomerase (TR) (Fajkus et al. 2019; Song et al. 2021). C) TRBs are associated with short telomeric sequences (*telo*-boxes) in the promoters of various genes in vivo, mainly with translation machinery genes (Schrumpfová et al. 2016). ORF, Open reading frame. D) *Telo*-box motifs recruit Polycomb repressive complexes (PRC2) via interactions of PRC2 subunits with TRB (Zhou et al. 2016; Zhou et al. 2018, this study) CLF, CURLY LEAF; SWN, SWINGER; EMF2, EMBRYONIC FLOWER 2; VRN2, VERNALIZATION 2. E) Histone H1 prevents the invasion of H3K27me3 and TRB1 over telomeres and long interstitial telomeric regions (Teano et al. 2020). F) TRB proteins, as subunits of the PEAT (PWO-EPCR-ARID-TRB) complex, are involved in heterochromatin formation and gene repression, but also have a locus-specific activating role, possibly through the promotion of histone acetylation (Tan et al. 2018; Tsuzuki and Wierzbicki 2018; Mikulski et al. 2019).

## Discussion

Although the TRBs were originally characterized as being associated with long arrays of telomeric repeats (Schrumpfová et al. 2004; Mozgová et al. 2008; Dvořáčková et al. 2010; Schrumpfová et al. 2014; Dreissig et al. 2017) (Fig. 8A), recent observations indicate broad engagement of TRB proteins in various cellular pathways. The most important TRB functions (Fig. 8 B-F) include interactions with the telomerase complex (Schrumpfová et al. 2014; Schořová et al. 2019), association with *telo*-boxes in the promoters mainly of translation machinery genes (Schrumpfová et al. 2016), recruitment of PRC2 and PEAT complexes to *telo*-boxes (Zhou et al. 2018; Tan et al. 2018; Tsuzuki and Wierzbicki 2018; Mikulski et al. 2019) or antagonisms between TRB1 and H1 at long interstitial telomeric DNA repeats (Teano et al. 2020).

### TRB4-5 are evolutionarily closer to TRBs from lower plants

Our phylogeny indicates that TRB proteins with N-terminal Myb-domains are conserved in plants and probably arose in Streptophyte algae within the family *Klebsormidiophyceae*. This corresponds to the transition of plants to a terrestrial habitat 800 Mya ago (Cheng et al. 2019) (Fig. 2). However, other groups of Streptophyte algae do not have TRB proteins, favoring a birth and death model of gene evolution or total divergence, such as to the TRF-like (TRFL) genes with a C-terminal Myb-domain.

Recurrent gene duplications over many generations have created orthologs and paralogs in the plant genome, giving rise to several new protein functions. In green algae, the majority of gene families contain only one gene (Clark and Donoghue 2018; Qiao et al. 2019), consistent with this, only one copy of the TRB protein was detected in Streptophyte algae. An increased number of TRB homologs were found in Embryophyta, which could be a result of WGD within these derived groups. In Mosses (taxon Bryophyta), the genome was duplicated in several independent events. Two WGD events in *Sphagnum* and *Physcomitrium* around 200 Mya and 120 Mya, respectively (Lang et al. 2018; Clark and Donoghue 2018), resulted in three almost identical TRB proteins clustered in the lineage TRB_A. The moss *Ceratodon* also underwent WGD, but unlike *Sphagnum* and *Physcomitrium*, the TRB duplication was not detected. No TRBs were found in hornworts and liverworts, but this may be due to the limited data available on these genomes.

Our results suggest diversification of the TRB gene family in seed plants which is linked with multiple subsequent or independent events of WGD. Lineage TRB_B seems to have evolved in the ancient past and is closely related to TRB_A in Bryophytes. The sub-lineage TRB_D is present in dicots in many families and embedded within the TRB_C lineage.

Within *Brassicaceae*, nested WGDs resulted in multiple TRB homologs. *A. thaliana* has five TRBs with closely related proteins TRB2-3 and TRB4-5. However, within Brassicales the situation differs. The genome of *Carica papaya* (*Caricaceae*) possesses only two TRB proteins, each belonging to one of the lineages TRB_B or TRB_C lineages. This observation is in agreement with an older paleoploidy event in the *Brassicaceae* lineage (β) that is not shared by *C. papaya* but is shared by all other analyzed *Brassicaceae* species (Dassanayake et al. 2011; Wang et al. 2011). As summarized by Rockinger et al. 2016, the ancestor of all *Caricaceae* underwent only a single WGD event, in comparison to the ancestor of *A. thaliana*, which underwent a more recent, additional round of WGD (α) (Bowers et al. 2003; Kagale et al. 2014). Similarly, in monocots, where several independent WGDs have occurred, an increased number of TRB proteins were found, e.g., six TRBs were detected in *Zea mays* (Figure 2B).

The complexity of plant genomes is extremely high, and annotations of plant genomes are undergoing intensive improvement at present (Kress et al. 2022). Further revision and reinvestigation of the TRB evolutionary tree may help to elucidate species-specific TRB variants such as the one identified in *Malus domestica*.

### Conserved structure of individual domains

Although TRB4-5 are grouped in a monophyletic lineage that is distant to TRB1-3 proteins, our assessment of predicted structural models implies that all TRB family members are folded into similar three-dimensional structures. The TRB Myb-like domain is very closely related to that of other telomere-binding proteins, including human TRF1 and TRF2 (Chong et al. 1995; van Steensel et al. 1998; Smogorzewska and de Lange 2004). TRF1 and TRF2 bind to DNA as preformed homodimers (Bianchi et al. 1997; Broccoli et al. 1997; Bianchi et al. 1999). It was suggested that in vitro the Myb-like binding domain of TRF1 binds to DNA essentially independently of the rest of the protein (König et al. 1998; Bianchi et al. 1999; Court et al. 2005). Specificity in DNA recognition of TRF1 and TRF2 is achieved by several direct contacts from aa side chains to the DNA, mainly via helix 3 and an extended N-terminal arm (Hanaoka et al. 2005; Court et al. 2005). Predicted models of the plant Myb-like domain (Fig. 1) show that the overall 3D structure is preserved in both plant and animal kingdoms, and that the surface mediating protein-DNA interactions are fully conserved. However, the surface side and extended N-terminal arm of the Myb-domain of proteins from the TRB_B family differ to those of the TRB_A or TRB_C families (Fig. 3).

Consistent with the conserved features of the Myb domain, our EMSA results showed the conserved ability of TRB4-5 proteins to bind telomeric sequences. Notably, not all proteins with Myb-like domains are able to bind telomeric repeats, as *Arabidopsis* proteins from the TRFL family, with this motif at the C-terminus, need an accessory Myb-extension domain for telomeric dsDNA interactions in vitro (Karamysheva et al. 2004; Ko et al. 2008). Moreover, unlike TRBs, even *Arabidopsis* plants deficient for all six TRFL proteins did not exhibit changes in telomere length or phenotypes associated with telomere dysfunction (Fulcher and Riha 2016). Our EMSA results showed the ability of TRB4-5 proteins to bind longer telomeric tracts as well as short *telo*-boxes, containing roughly one telomeric repeat flanked by non-telomeric sequence (Fig. 4). These observations suggest that TRB4-5 proteins may be associated with cis-regulatory elements in the promoter regions of *Arabidopsis* genes as was described for TRB1-3 (Zhou et al. 2016; Schrumpfová et al. 2016). The presence of only one telomeric repeat in the *telo*-box raises the question of whether TRB proteins operate on promoter regions as monomers/dimers or multimers. Based on our quantitative DNA-binding study, we propose that TRB1 and TRB3 bind long telomeric DNA arrays with the stoichiometry of one protein monomer per one telomeric repeat (Hofr et al. 2009), but the stoichiometry of TRBs in regulatory complexes associated with *telo*-boxes needs further elucidation.

The sequence-specific interaction between telomeric dsDNA and TRB1-3 is mediated predominantly by the Myb-like domain, although additional domains from TRBs can also contribute to non-specific DNA interactions (Schrumpfová et al. 2004; Mozgová et al. 2008; Hofr et al. 2009). The H1/5-like domains of TRB proteins belong to the same group as central globular domain of core H1 histones, the incorporation of which directly influences the physicochemical properties of the chromatin fiber and further modulates nucleosome distribution, chromatin compaction and contributes to the local variation in transcriptional activity by affecting the accessibility of transcription factors and RNA polymerases to chromatin (Fan and Roberts 2006; Zhou et al. 2015; Hergeth and Schneider 2015; Bednar et al. 2017; Fyodorov et al. 2018). The H1/5-like domain of TRB mediates non-specific DNA interactions (Mozgová et al. 2008) as well as interactions with the other members of TRB family and also with the POT1b protein (Schrumpfová et al. 2008). The globular domain of H1 adopts a winged-helix fold with a “wing” defined by two β-sheets. 3D model predictions suggest only minor differences between TRB4-5 and TRB1-3 within the H1/5-like domain, as the loop between two antiparallel β-sheets in the H1/5-like domain of TRB4-5 is longer than in TRB1-3.

Coiled-coil domains are structural motifs that consist of two or more α-helical peptides that are wrapped around each other in a superhelical fashion that may mediate interactions between proteins (Lupas and Gruber 2005; Apostolovic et al. 2010). Only minor differences were predicted for the 3D structure of the coiled coil domain of TRB4-5 and TRB1-3. However, the motifs at the C-terminal sections of the coiled-coil domains in TRBs from TRB_B differ from those motifs at the C-terminal parts of TRB_C lineages, and clearly distinguish these two evolutionarily distinct groups.

### TRB4-5 differ in subcellular localization from TRB1-3

In previous studies, it was shown that TRB1-3 are highly dynamic DNA-binding proteins with cell-cycle regulated localization. During interphase, GFP-fused TRB1-3 proteins are preferentially localized in the nucleus, with a strong nucleolar signal and relatively strong nuclear speckles of different sizes (Dvořáčková et al. 2010; Schrumpfová et al. 2014; Zhou et al. 2018). A similar pattern was observed in transiently or stably transformed *Arabidopsis* cells (Dvořáčková et al. 2010) or in tobacco epidermal cells (Schrumpfová et al. 2014) despite the fact that tobacco telomeres are dispersed throughout the nucleus (so-called non-Rabl chromosome configuration), while *Arabidopsis* telomeres are clustered around the nucleolus (rosette-like chromosome configuration) (Shan et al. 2021). Our results show that even native TRBs in isolated *Arabidopsis* nuclei can be detected with a specific antibody, and are localized in the speckles. Some of these speckles in the vicinity of the *Arabidopsis* nucleoli might be telomeric (Dvořáčková et al. 2010), or might be Cajal bodies (Dvořáčková 2010), as was demonstrated for speckles detected in tobacco nuclei.

Here we show that TRB4 and TRB5 fused to GFP have, in contrast to TRB1-3, a distinct subcellular localization pattern (Fig. 5B, C). In addition, unlike all other members of the TRB family, TRB5 is preferentially localized in the cytoplasm. It remains to be clarified whether it is sequestered there or plays a specific functional role.

Our Y2H or BiFC assays proved that TRB4-5 could form homo- and hetero-dimers, as observed in TRB1-3 (Kuchař and Fajkus 2004; Schrumpfová et al. 2004). In addition to dimers, TRB1-3 are capable of forming both homo- and heterotypic multimers via their H1/5-like domain (Schrumpfová et al. 2004; Mozgová et al. 2008; Hofr et al. 2009). Similar multimerization can also be assumed for TRB4 and TRB5, as we observed the formation of high molecular weight complexes for TRB4-5 in EMSAs, which did not migrate into the gel (Warren et al. 2003).

Despite distinct subcellular localization of GFP-TRB4 and GFP-TRB5, we observed mutual interactions between all TRB family members using BiFC, predominantly in the nuclear speckles (Fig. 6B). Only the TRB5 homodimeric interaction was found to be clearly cytoplasmic with nuclear speckles of decreased size compared to the size of speckles in the other homodimeric or mutual TRB interactions. Our BiFC assays showed that TRB1-3, but not TRB4, have the ability to drag TRB5 into the nucleolus, as TRB4 does not interact with TRB5 in the nucleolus. We can assume that TRB proteins form various heteromers in different subcellular compartments that might possess different functions related to distinct biochemical pathways.

### Interconnection of TRBs with various protein complexes

Even though TRB4-5 show slightly distinct localization patterns compared to TRB1-3, it appears that all TRBs may interact with similar partners in vitro (Fig. 7). TRB4-5 proteins directly interact with the N-terminal domains of TERT as was also shown for TRB1-3 proteins (Schrumpfová et al. 2014). Additionally, TRB4-5 interact with both RUVBL1 and RUVBL2A proteins, which may imply that they may mediate interactions in the trimeric complex RUVBL-TRB-TERT as was proposed for TRB1-3 (Schořová et al. 2019). Telomerase might also be modulated by POT1 proteins: while *A. thaliana* POT1a positively regulates telomerase activity, whereas POT1b is proposed to negatively regulate telomerase and promote chromosome end protection (Beilstein et al. 2015). We observed not only the expected interaction between TRB4-5 and POT1b (Schrumpfová et al. 2008), but also revealed novel interactions of TRB proteins with POT1a. Altogether, we can assume that TRB4-5 are associated with the telomerase complex as was proved for TRB1-3.

In addition to telomeres (Schrumpfová et al. 2014), TRB1-3 regulate the PRC2 target genes (Zhou et al. 2016; Zhou et al. 2018). Interestingly, H3K27me3 is highly abundant at telomeres, including those of humans (Montero et al. 2018) and *Arabidopsis* (Vaquero-Sedas et al. 2012; Adamusová et al. 2020). The observation that not only TRB1-3 (Zhou et al. 2018), but also TRB4-5, physically interact with homologs of E(z) subunit of PRC2 complex named CLF and SWN suggests that all TRBs can target PRC2 to Polycomb response elements (PREs) including *telo*-boxes (Deng et al. 2013; Zhou et al. 2016; Godwin and Farrona 2022). Moreover, the novel interaction between TRBs and Su(z)12 *A. thaliana* homologues EMF2 and VRN2 described here, tightly interconnects all TRBs with the core PRC2 components. These observations support the recently published observation that *O. sativa* single Myb transcription factor TRBF2 forms phase-separated droplets, which aggregate with PRC2 via rice OsCLF and OsEMF2 (Xuan et al. 2022).

In *Arabidopsis*, the PWO1 protein interacts with all three E(z) homologs, including CLF and SWN through its conserved N-terminal PWWP domain (Mikulski et al. 2019). Furthermore, PWO1-3 are associated with members of the PEAT complex, which was recently identified as being able to silence transposable elements (Tan et al. 2018). Our data suggest that the interaction between TRBs and PWOs is not restricted to only TRB1 and 2 (Tan et al. 2018), instead other TRB members, including TRB4-5, interact with PWWP domains located at the the N-terminus of PWO1 and PWO3 from *A. thaliana*. Additionally, the PWWP domain of PWO2 is recognized by *Arabidopsis* TRB4. Interestingly, in tobacco PWO1 tethers CLF to nuclear speckles (Hohenstatt et al. 2018; Mikulski et al. 2019) which have a similar distribution to speckles of TRB proteins.

Overall, we conclude that TRB proteins, including the newly characterized TRB4-5, are associated with several complexes, including telomerase, PRC2 or PEAT complexes. However, TRBs are unlikely to be permanently associated with all of these complexes (Tan et al. 2018; Schubert 2019), and we might speculate that TRB subunits are partially interchangeable within these complexes.

It should be noted that the number of identified interaction partners of TRBs in *Arabidopsis* may increase in the future, as these promiscuous proteins may play a role in various additional biochemical processes that are not yet elucidated. For example, the *A. thaliana TRB1* gene is responsive to several types of hormones, including jasmonate (JA) (Yanhui et al. 2006); TRB homologue from apple dynamically modulates JA-mediated accumulation of anthocyanin and proanthocyanidin (An et al. 2021); soybean TRB homologue was identified as candidate gene regulating total soluble sugar in soybean seeds (Xu et al. 2022).

## CONCLUSION

Proteins from the TRB family are plant-specific and apparently first evolved in lower plants. We speculate, that due to WGDs one ancestral TRB was multiplied to the current five TRB members in *A. thaliana* increasing the potential of diversification of their particular functions. Their Myb-like domain specifically targets these proteins to telomeric sequences located terminally (Schrumpfová et al. 2004; Mozgová et al. 2008; Dvořáčková et al. 2010; Schrumpfová et al. 2014; Dreissig et al. 2017) or interstitially (Schrumpfová et al. 2016). Additionally, the versatile interactions of TRBs with other proteins contribute to the multiple functions that they adopt in the cell nucleus, including participation in telomerase biogenesis (Schrumpfová et al. 2014; Schořová et al. 2019), recruitment of PRC2 or PEAT complexes (Zhou et al. 2016; Zhou et al. 2018; Tan et al. 2018; Tsuzuki and Wierzbicki 2018; Mikulski et al. 2019) or competing with H1 for binding to interstitially localized telomeric sequences (Teano et al. 2020). The cytoplasmic localization of TRB5 and its implications deserve further investigation. As additional functions and interaction partners of TRBs are discovered, it can be expected that research in plants will lead to a better understanding of the mode of action of the different TRBs and also to the elucidation of novel functions.

## Materials and methods

### Primers

The sequences of all primers and probes used in this study are provided in Supplemental Table 4.

### Plant material

For transient assays, *Nicotiana benthamiana* plants were grown in soil in LD conditions up to 4 weeks and subsequently used for *Agrobacterium tumefaciens* infiltration.

### Phylogenetic analyses

We combined two homology searches based on *A. thaliana* TRB1-5. First, we searched completely sequenced genomes using Phytozome v12 and second using BLASTP from available databases (NCBI) and publicly available sequences.

Protein sequences were aligned using the Clustal Omega (Sievers et al. 2011) algorithm in the Mobyle platform (Néron et al. 2009), with homology detection by HMM–HMM comparisons. We screened data after alignment in the BioEdit program (Hall 1999).

Maximum likelihood (ML) analyses of the matrices were performed in RAxML 8.2.4 (Stamatakis 2014) to examine differences in optimality between alternative topologies. 1000 replications were run for bootstrap values. Phylogenetic trees were constructed and modified with iTOL v3.4 (Letunic and Bork 2016). The MEME search was set to identify domains and conserved amino acid (aa) sequence motifs under these conditions: a maximum of 15 motifs for each protein with a wide sequence motif from 2 to 50 and a total number of sites from 2 to 600 MEME 4.11.2 (Bailey et al. 2009). The evolutionarily conserved aa residues were visualized using ConSurf 2016 (Ashkenazy et al. 2016).

### Analysis of protein structures

The AlphaFold (Jumper et al, 2021; Varadi et al, 2022) and SwissModel (Waterhouse et al, 2018) tools were used to generate in silico protein models. Structural models were compared as previously described (Palecek & Gruber, 2015). All structures including electrostatic potential of their molecular surfaces were visualized using PyMOL Molecular Graphics System, Version 2.4.1, Schrödinger, LLC.

### Cloning

For yeast two-hybrid assays (Y2H), most of the Y2H constructs in pGADT7-DEST or pGBKT7-DEST (Horák et al. 2008) were prepared previously: TERT constructs (RID1, TEN, Fw1N, Fw3_NLS, Fw3N and RT) were reported in Majerská et al. 2017, TRB1-3 were reported in Schrumpfová et al. 2014, RUVBL1 and RUVBL2A were reported in Schořová et al. 2019, POT1a and POT1b were reported in Majerská et al. 2017, SWN_Δ_SET and CLF_Δ_SET were reported in Chanvivattana et al. 2004 and Hohenstatt et al. 2018). PWO1 was reported in Hohenstatt et al. 2018, VRN2 and EMF2 were reported in Lindner et al. 2013. The coding sequences of PWO2 and the fragment of PWO3 were cloned in the pGBKT7 and pGADT7 vectors (Clontech), passing through pDONR221 (Invitrogen) as described in Hohenstatt et al. 2018.

For the TRB4 construct, the cloned cDNA sequence of *TRB4* (G60951 from Arabidopsis Biological Resource Center; ABRC, https://abrc.osu.edu/) in pENTR223 was used as the entry clone. Site-directed mutagenesis using the QuikChange XL Site-Directed Mutagenesis Kit (Agilent Technologies) was performed following the manufacturer’s instructions to remove the mutation V122A in the protein encoded by pENTR223-TRB4 as described in Wiese et al. 2021. To generate an entry vector containing the cDNA sequence of the *TRB5* gene, the total RNA from 7-day-old *A. thaliana* Col-0 seedlings isolated by TRI reagent (Molecular Research Center) was used for cDNA preparation using M-MuLV Reverse Transcriptase (New England Biolabs). The cDNA sequence of *TRB5* was amplified using gene specific Gateway-compatible primers according to the manufacturer’s instructions with primers specified in Supplemental Table 4, and the RT-PCR products were recombined into the Gateway donor vector pDONR207 (Invitrogen). DNA fragments were introduced into the destination Gateway vectors pGBKT7-GW, pGADT7-GW (Addgene) using the LR recombinase reaction (Invitrogen).

For BiFC experiments, the Multisite Gateway^®^ system (Invitrogen) was used to create pBiFCt-2in1 constructs (Grefen and Blatt 2012). The genes encoding *TRB1-5* from *A. thaliana* Col-0 were PCR-amplified from the constructs used in the Y2H system described above by two-step PCR using primers specified in Supplemental Table 4 and Phusion™ High-Fidelity DNA Polymerase (Thermo Fisher Scientific) as described in Wiese et al. 2021. The amplicons with *TRB1,2,3,5* genes were cloned into Gateway pDONR221 entry vectors (Thermo Fisher Scientific) carrying either attP1-P4 or attP3-P2 recombination sites using the BP Clonase™ II enzyme mix (Thermo Fisher Scientific). To generate pDONR221 entry clones carrying the *TRB4* gene, the In-Fusion® Snap Assembly cloning kit (Takara Bio USA) was used, where PCR-generated TRB4 amplicons with one half of att-sites obtained in the first Phusion PCR step of the Gateway® cloning were fused with linearized PCR-generated pDONR221 backbone with appropriate attL sites by recognizing 15-base pairs (bp) overlaps at their ends according to the manufacturer’s instructions. All entry clones were subsequently used in LR Clonase™ II Plus (Thermo Fisher Scientific) reactions to create pBiFCt-2in1-CC and pBiFCt-2in1-NN expression constructs harboring two protein coding regions C- or N-terminally fused to either the N- or C-terminal eYFP halves (e.g., TRB1-nYFP and TRB1-cYFP). After verification by Sanger sequencing (Macrogene), the constructs were used for transient expression in *N. benthamiana* leaves.

For transient expression of TRB1-5 fused with GFP in *N. benthamiana* leaf epidermal cells, the specific entry clones described above (TRB1-3, 5 in pDONR207 and TRB4 in pENTR223) were used in LR Clonase™ II (Thermo Fisher Scientific) reactions to create the pGWB6 (N-terminal GFP fusion under the 35S promoter) expression vectors (Nakagawa et al. 2007). To label nucleolus or nucleoplasm, we co-transfected *N. benthamiana* leaf epidermal cells with constructs expressing Fibrillarin1-mRFP (Koroleva et al. 2009) or SRp34-mRFP (Lorković et al. 2004; Koroleva et al. 2009), respectively.

For protein expression in *Escherichia coli*, constructs in pDONR207 (for TRB1 and TRB5) or in pENTR223 (for TRB4) were used as donor vectors for LR Clonase™ reactions (Thermo Fisher Scientific), where genes coding proteins of interest (TRB1, 4 and 5) were transferred into the destination vector pHGWA (Busso et al. 2005).

### Yeast two-hybrid assays

Y2H was performed using the Matchmaker TM GAL4-based two-hybrid system (Clontech) as described in Schrumpfová et al. 2014. Successful co-transformation of each bait/prey combination into *Saccharomyces cerevisiae* PJ69-4a was confirmed on SD plates lacking Leu and Trp, and interactions assessed on SD medium lacking Leu, Trp and His (with or without 1mM or 3mM 3-aminotriazol (3-AT)) or SD medium lacking Leu, Trp and Ade at 30 °C. Co-transformation with an empty vector and homodimerization of the TRB1 protein served as a negative and positive control, respectively (Schrumpfová et al. 2014). Each combination was co-transformed at least three times, and at least three independent drop tests were carried out. Protein expression was verified as described in Schořová et al. 2019.

### In vitro translation and co-immunoprecipitation

Proteins were expressed from constructs similar to those used in the yeast two-hybrid system with a hemagglutinin tag (pGADT7-DEST; VRN2, EMF2 and POT1a proteins) or a myc-tag (pGBKT7-DEST; TRB proteins) using a TNT Quick Coupled Transcription/Translation system (Promega) in 50 μl reaction volumes according to the manufacturer’s instructions. The VRN2, EMF2 or POT1a proteins were radioactively labelled using ^35^S-Met. The co-immunoprecipitation procedure was performed as described by Schrumpfová et al., 2008. Input, Unbound and Bound fractions were separated by 10% SDS – PAGE, and analyzed using a FLA7000 imager (Fuji-film).

### Transient heterologous expression

*A. tumefaciens* competent cells (strain GV3101) were transformed with selected expression clones and selected on YEB medium supplemented with gentamycin (50 μg/mL), rifampicin (50 μg/mL), and a vector-specific selection agent (pBiFCt-2in1: spectinomycin 100 μg/mL, pGWB6: kanamycin and hygromycin both 50 μg/mL, pROK2: kanamycin 50 μg/mL) at 28 °C for 48 h. Colonies were inoculated in the same media lacking agar and grown overnight at 28 °C. Bacterial cells of overnight cultures were pelleted by centrifugation (5 min at 1620 g), washed twice, re-suspended, and diluted to an OD_600_ of 0.5 with infiltration medium (10 mM MES pH 5.6, 10 mM MgCl_2_ and 200 μM acetosyringone). A suspension of *Agrobacterium* cells carrying the p19 repressor plasmid was added in a 1:1 ratio with *Agrobacterium* suspensions harboring plasmids of interest to suppress gene silencing and enhance transient expression (Gehl et al. 2009). Mixed suspensions were incubated with moderate shaking for 1.5-2 h at room temperature and subsequently injected into the abaxial side of leaves of 4-week-old *N. benthamiana* plant. On the third day after infiltration, tobacco epidermal cells were microscopically analyzed. Microscopy images were acquired using the Zeiss LSM880 laser scanning microscope (Axio Observer Z1, inverted) with Definite Focus 2 (excitation 488 nm for GFP/YFP and 561 nm for mRFP). Images were processed using Fiji/ImageJ (Schindelin et al. 2012) and Adobe Photoshop CS6 (Adobe, San Jose, CA, USA) software.

### Nuclei isolation and immunofluorescence

Isolation of nuclei was performed using 10-day-old seedlings as described in (McKeown et al. 2008). Nuclei were centrifuged for 20 min at 350 g and 4 °C, resuspended in 1× PBS and spotted onto slides. Nuclei were briefly dried at 4 °C, then fixed in 4% paraformaldehyde in 1× PBS, 0.5% Triton-X for 10 min. Nuclei were rinsed three times in 1x PBS, blocked in 5% goat serum with 0.05% Tween-20 for 30 min at RT and incubated overnight with antibody anti-TRB 5.2 (Schrumpfová et al. 2014) diluted 1:300 in 5% BSA in 1× PBS. Nuclei were washed three times for 5 min in 1× PBS supplemented with 0,05% Tween-20 (PBST), then incubated 1 h with an anti-mouse Alexa 488 antibody, Invitrogen (1:750 dilution). Slides were washed three times for 5 min in 1× PBST, then dehydrated in ethanol series as described in Kutashev et al. 2021. Coverslips were mounted in 4′,6-Diamidine-2′-phenylindole dihydrochloride (DAPI), 2 μg/mL in Vectashield and imaged using fluorescence using a Zeiss AxioImager Z1 epifluorescence microscope.

### Expression of TRB1, 4 and 5 in *E. coli*

Proteins fused with His-tags were expressed in *E. coli* (BL21(DE3) RIPL) from the destination vector pHGWA. BL21(DE3) RIPL were grown in Luria-Bertani medium suplemented with ampicilin and chloramphenicol to final concentration 100ug/ml and 12,5ug/ml at 37 °C until OD600 reached 0.5. Cells were cultured for 4 h at 25 °C after the addition of IPTG to a final concentration of 0.5 mM. Cells were collected by centrifugation (8,000 g, 8 min, 4 °C). The cell pellet was dissolved in lysis buffer containing 50 mM sodium phosphate (pH 8.0), 300 mM NaCl, 10 mM imidazole, 5mM β-mercaptoethanol, and protease inhibitor cocktail cOmplete tablets EDTA-free (Roche). The cell suspension was sonicated on ice for 6 min of process time with 1 s pulse and 2 s pause (Misonix). Cell lysate supernatant was collected after centrifugation at 20,000 g, 4 °C for 1 h. Proteins were purified by immobilized-metal affinity chromatography using TALON® metal affinity resin (Clontech), where filtered supernatant (0.45 μm filter) was mixed with TALON® beads and incubated for 1 hour. The proteins of interest were eluted with 100 mM imidazole in the same buffer. These proteins were then concentrated, and the buffer was exchanged for 50 mM sodium phosphate (pH 7.0), 300 mM NaCl by ultrafiltration (Amicon 3 K/30 K, Millipore). The concentration of purified proteins was determined using the Bradford assay. We evaluated protein purity using SDS-polyacrylamide gel electrophoresis, with gels stained by Bio-Safe Coomassie G250 (Bio-Rad). To ensure successful purification, the purified proteins were detected by specific monoclonal mouse anti-polyhistidine antibody as described previously (Schrumpfová et al. 2004) and by matrix-assisted laser desorption-ionization tandem mass spectrometry (MALDI-MS/MS) (CEITEC Proteomics Core Facility).

### Electrophoresis mobility shift assay (EMSA)

DNA probes and competitors used are described in Supplemental Table 4. To reduce a non-specific DNA–protein binding, 10 pmol of purified TRB1, 4 or 5 were preincubated with 1, 10, or 100 pmol of a specific competitor (oligodeoxynucleotides of non-telomeric or competitor telomeric sequence in double-stranded form, as indicated in Results) for 20 min in EMSA buffer (10 mM Tris–HCl pH 8.0, 1 mM EDTA, 1 mM dithiothreitol, 50 mM NaCl, and 5% w/v glycerol). Probes were end-labelled using [γ-^32^P]ATP and polynucleotide kinase (New England Biolabs), according to (Sambrook et al. 1989). A probe (1 pmol) was added to the reaction mixture on ice and incubated for 20 min before the mixture was loaded onto a 7.5% w/v non-denaturing polyacrylamide gel (AA:BIS = 37.5:1, 0.5× TBE, 1.5 mm thick). Electrophoresis was at 15 °C for 3 h at 10 V/cm in a precooled buffer. Signals of labelled oligonucleotides were detected using a phosphorimager FLA-7000IP (Fujifilm).

## Author contribution

Petra Prochazkova Schrumpfova and David Honys designed the study, supervised the project. Petra Prochazkova Schrumpfova wrote the manuscript with support from david Honys, Jan Palecek, Daniel Schubert. Lenka Steinbachova, Zuzana Gadiou, Alzbeta Kusova, Tereza Prerovska, Claire Jourdain and Tino Stricker performed cloning, Lenka Steinbachova performed localization and BiFC, Alzbeta Kusova. and Ahmed Khan performed Y2H, tereza Prerovska performed EMSA, Gabriela Rigoova performed Co-IP, Jan Palecek analyzed 3D structures, Lenka Zaveska Drabkova performed evolution analysis. Petra Prochazkova Schrumpfova conducted experiments and analyzed data.

## ACCESSION NUMBERS

TRB1 (AT1G49950); TRB2, formerly TBP3 (AT5G67580); TRB3, formerly TBP2 (AT3G49850); TRB4 AT1G17520; TRB5 AT1G72740; RUVBL1 (AT5G22330); RUVBL2A (AT5G67630); TERT (AT5G16850); POT1a (AT2G05210); POT1b (AT5G06310); Fibrillarin1 (AT5G52470); SRp34 (AT1G02840); EMF2 (At5G51230); VRN2 (At4G16845); CLF (At2G23380); SWN (At4G02020); PWO1 also named as PWWP1 (At3G03140); PWO2 also named as PWWP2 (At1G51745); PWO3 also named as PWWP3 (At3G21295)

## Acknowledgments

This work was supported by the Czech Science Foundation projects 21-15841S (P.P.S., M.K., D.H.) and 20-01331X (A.K), and Czech Ministry of Education, Youth and Sports grant number LTAUSA18115 (L.S.). The Plant Sciences Core Facility of CEITEC Masaryk University is acknowledged for its technical support. We acknowledge the Imaging Facility of the Institute of Experimental Botany AS CR supported by the MEYS CR (LM2018129 Czech-BioImaging) and IEB AS CR.

We acknowledge Sara Farrona (Plant and AgriBiosciences Centre, National University Ireland, IR) for providing us with *SWN, CLF* and *EMF2* constructs. Moreover, we acknowledge Eva Sykorova (Institute of Biophysics of the Czech Academy of Sciences, Brno, Czech Republic) for providing us constructs with *POT1a-b* and *TERT* fragments and Ali Pendle (John Innes Centre, Norwich, United Kingdom) for the sub-nuclear localization markers. We also acknowledge Yvona Stražická and Michal Franek for the help with the immunofluorescence on *Arabidopsis* nuclei.

We acknowledge Leon Jenner and Iva Mozgová for proofreading and manuscript editing.

The access to the computing and storage facilities owned by parties and projects contributing to the National Grid Infrastructure MetaCentrum provided under the programme Projects of Large Infrastructure for Research, Development, and Innovations (LM2010005) was highly appreciated, as was the access to the CERIT-SC computing and storage facilities provided under the programme Center CERIT Scientific Cloud<u>°</u>, part of the Operational Program Research and Development for Innovations, reg. no. CZ.1.05/3.2.00/08.0144.

## Supplemental Figures

**Supplemental Fig. 1.**
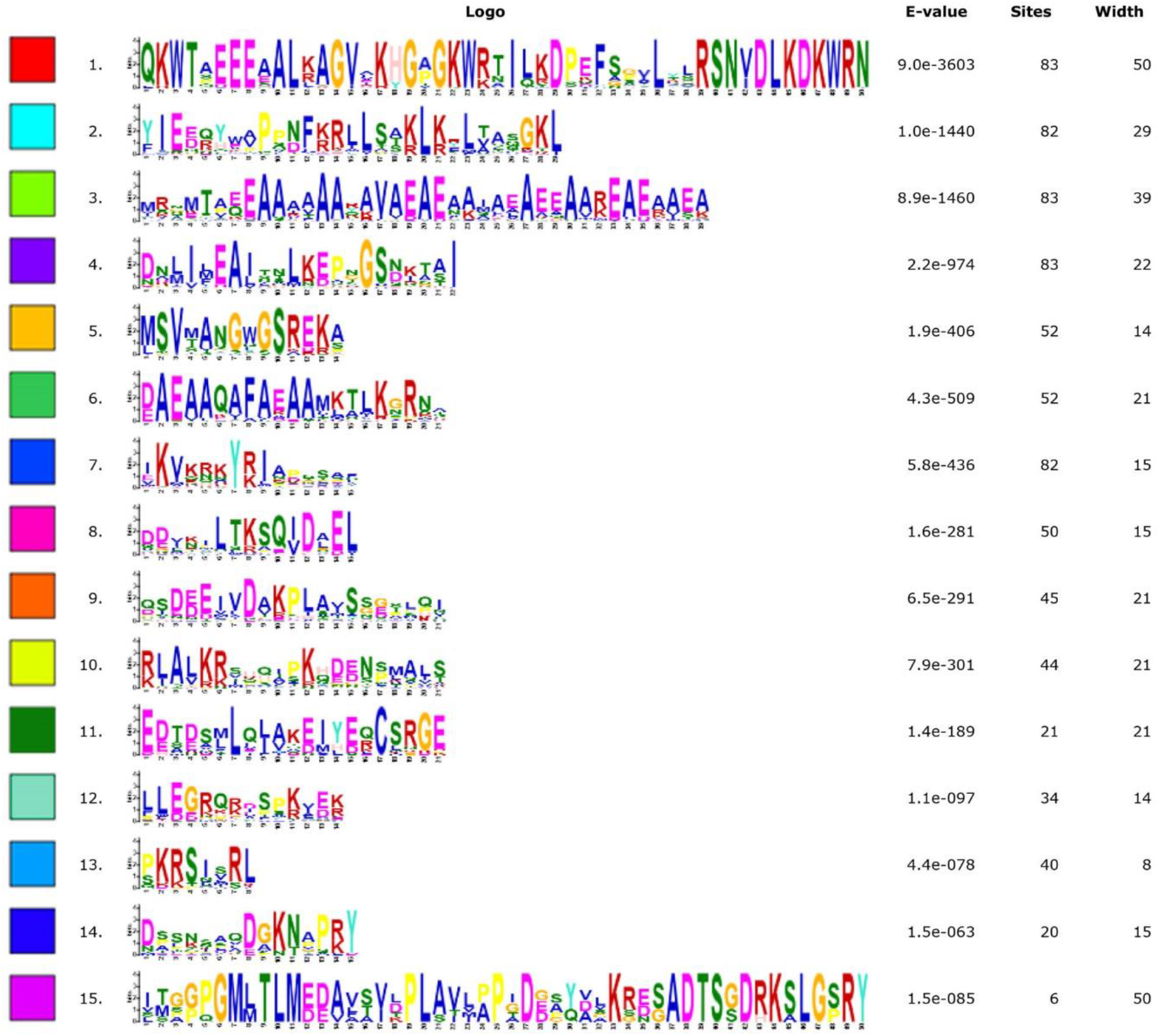
The 15 most conserved motifs in plant TRB protein motifs. Motifs are ranked and ordered by highest probability of occurrence. Motif 1 corresponds to the MYB-like domain. Motifs 2, 4 and 7 belong to the H1/H5-like domain. Motifs 3 and 6 or 3 and 11 create the coiled-coil domain.

**Supplemental Fig. 2.**
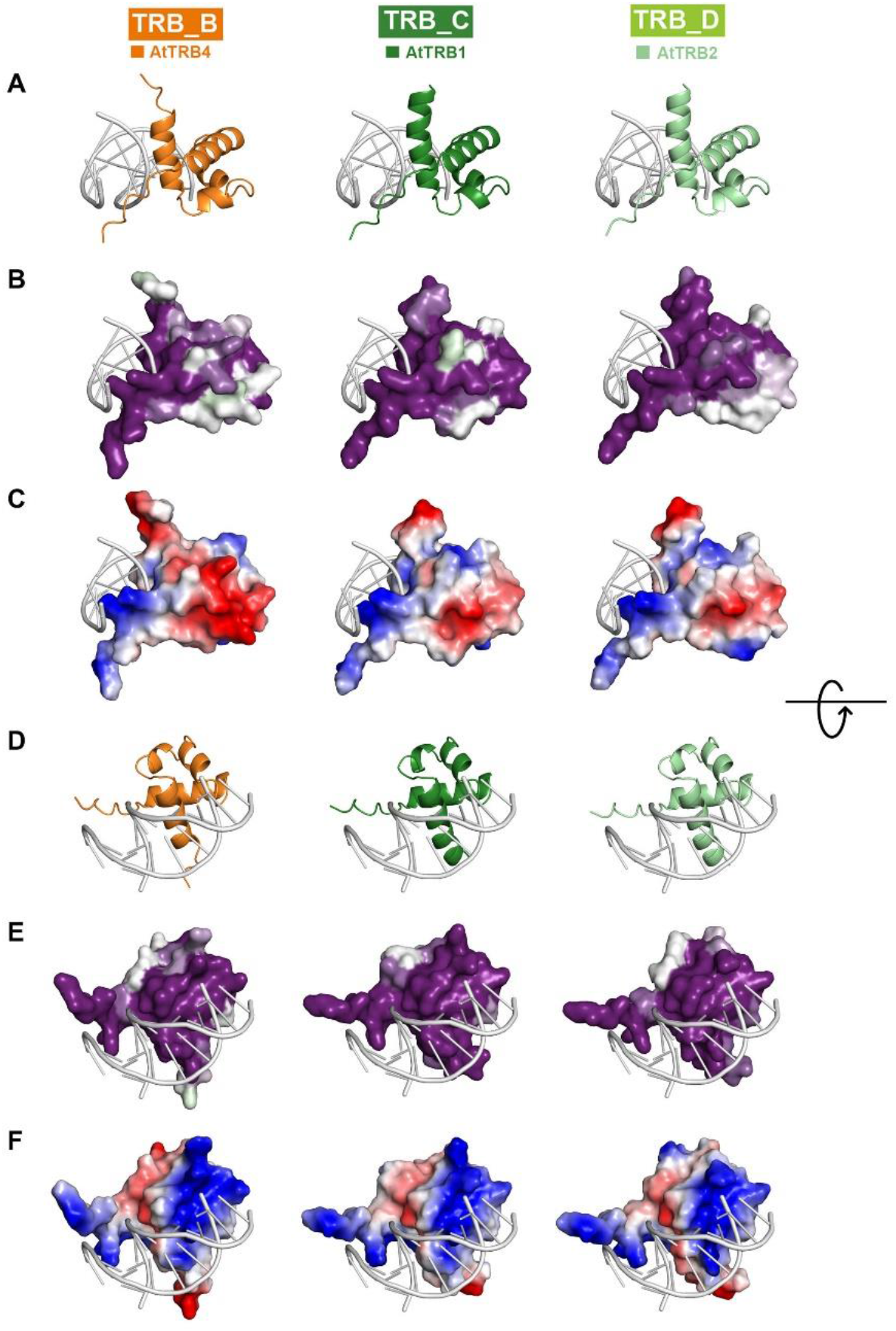
Conserved residues and electrostatic charge visualization of the Myb-like domain in *Arabidopsis* members of Dicots from TRB_B, TRB_C and TRB_D lineages. The representative members of Dicots from TRB_B, TRB_C and TRB_D lineages, namely *A. thaliana* TRB4, TRB1 and TRB2, respectively, were analyzed. The three-dimensional model of the Myb-like domains from the site opposing the DNA-binding (A-C) and from the DNA binding (D-F) viewpoints are based on the hTRF2-DNA interaction model (PDB: 1WOU) (Court et al. 2005). B) and E) The evolutionary dynamics of aa substitutions among aa residues were visualized in ConSurf 2016 (Ashkenazy 2016 doi.org/10.1093/nar/gkw408). The conservation of residues is presented in a scale, where the most conserved residues are shown in dark magenta and non-conserved residues as white. C) and F) Surface models showing the charge on the Myb-like domains. Residue charges are coded as red for acidic, blue for basic, and white for neutral, visualized using PyMol, Version 2.4.1, Schrödinger, LLC.

**Supplemental Fig. 3.**
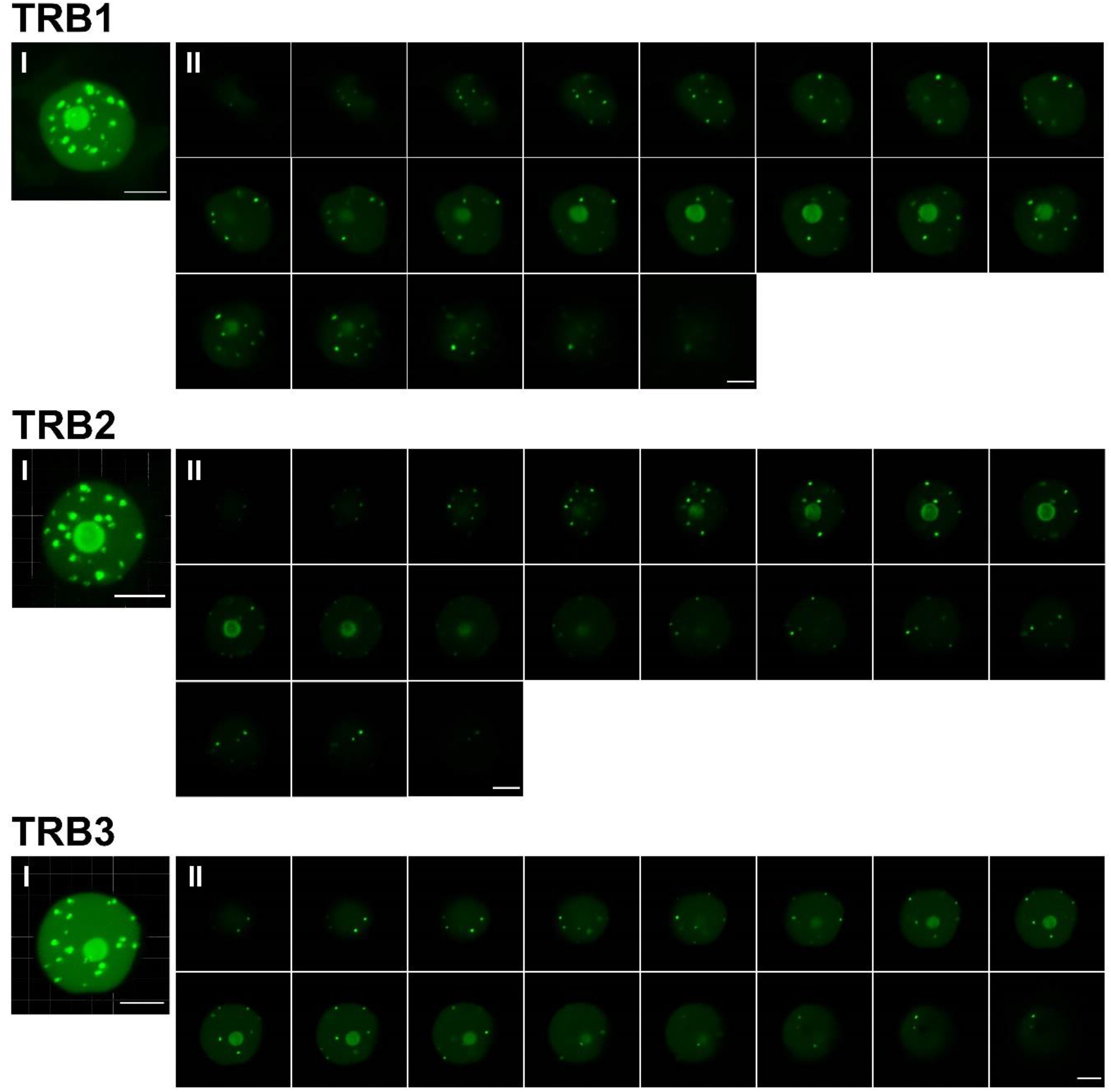
Maximum intensity projections and Z-stacks of TRB1-3 GFP-fusion proteins in nuclei of *N. benthamiana* leaf epidermal cells. TRB1-3 fused with GFP (N-terminal fusions), expressed in *N. benthamiana* leaf epidermal cells and observed by confocal microscopy. Figures represent Maximum Intensity projections (I) of entire Z-stack images of nuclei (II; TRB1 - 0,53 μm each; TRB2 and TRB3 – 0,47 μm each). Scale bars = 5 μm.

**Supplemental Fig. 4.**
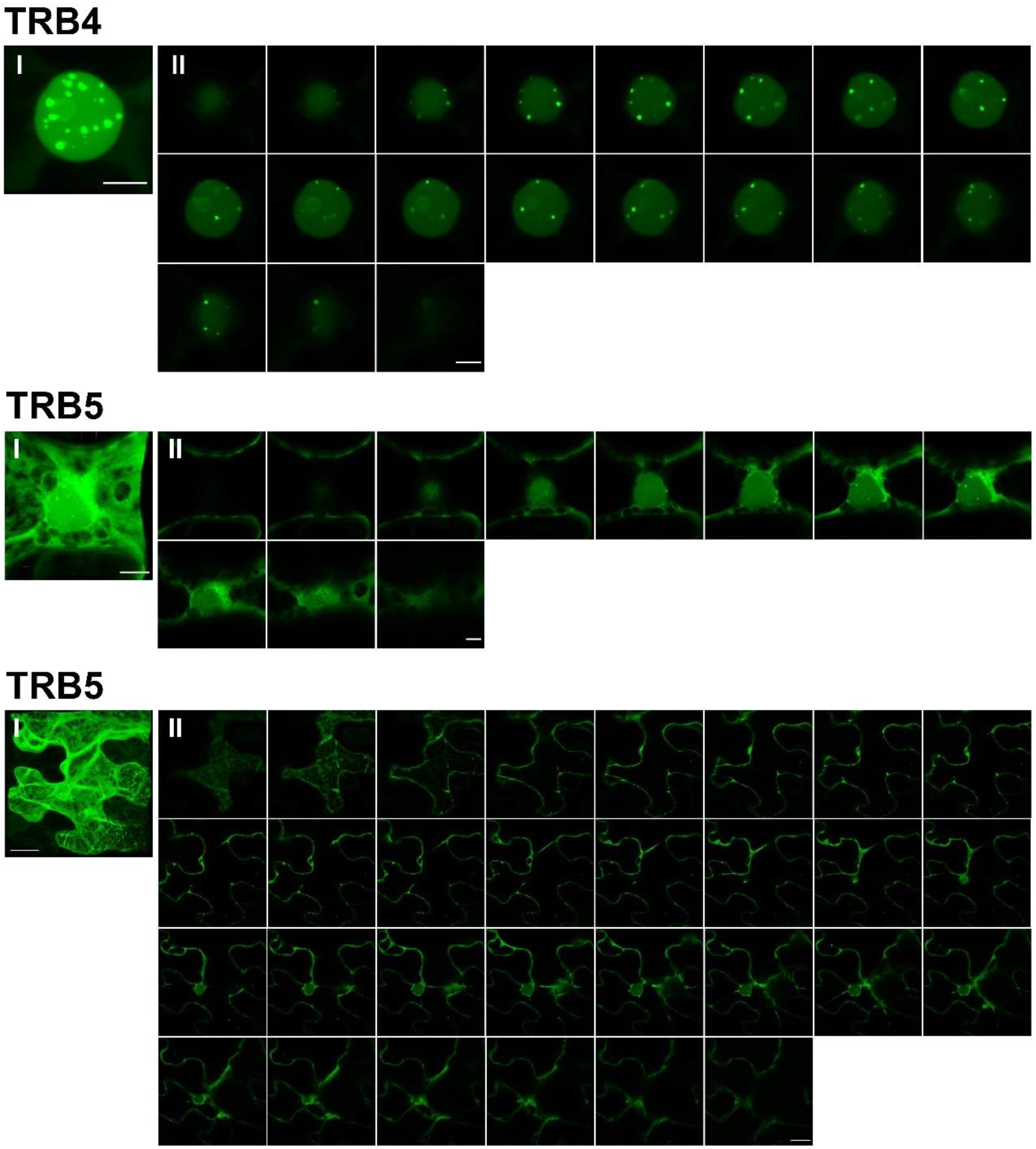
Maximum intensity projections and Z-stacks of TRB4-5 GFP-fusion proteins in *N. benthamiana* leaf epidermal cells. TRB4-5 fused with GFP (N-terminal fusions), expressed in *N. benthamiana* epidermal cells, and observed by confocal microscopy. Figures represent Maximum Intensity projections (I) of entire Z-stack images (II) of nuclei (TRB4 - 0,52 μm each; TRB5 - 0,82 μm each; scale bars = 5 μm) and whole epidermal cell (TRB5 - 0,78 μm each; scale bar = 20 μm).

**Supplemental Fig. 5.**
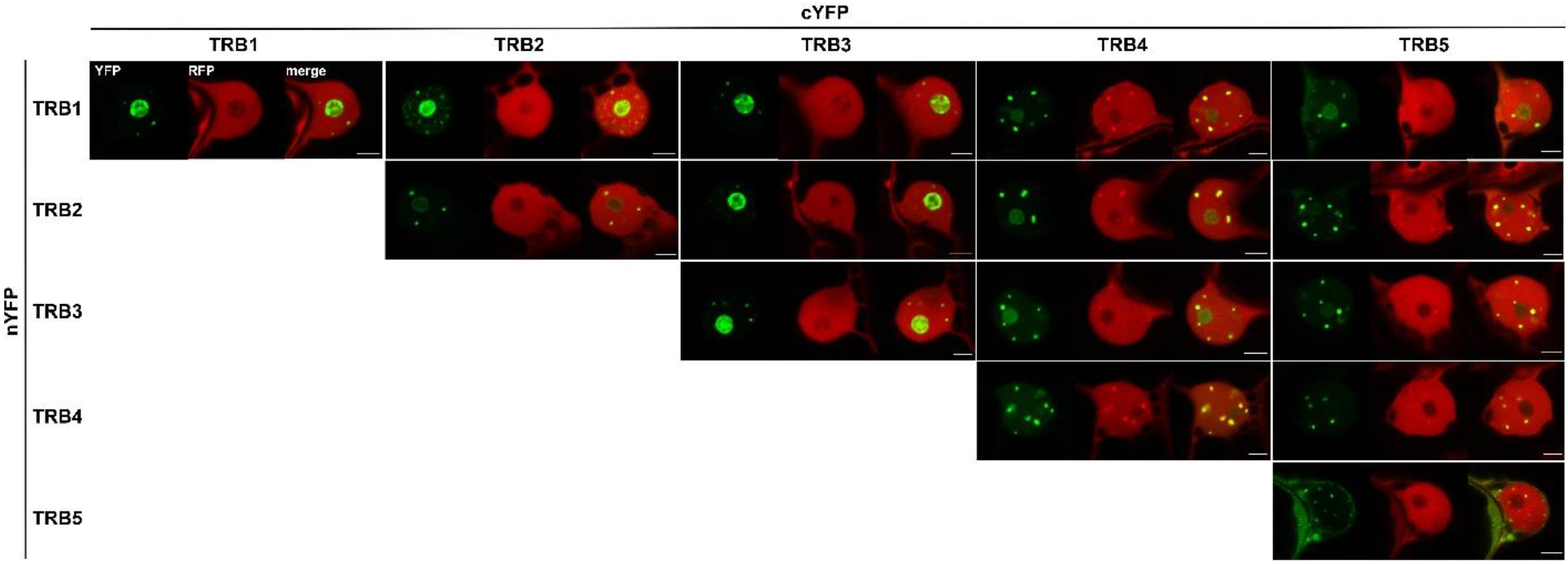
Mutual TRB interactions detected by BiFC in *N. benthamiana*. Protein-protein interactions of TRB proteins fused with nYFP or cYFP part were detected in *N. benthamiana* leaf epidermal cells by confocal microscopy. Shown here are single images of fluorescence signals from individual channels *(YFP*, Yellow fluorescence protein; *RFP*, red fluorescence protein – an internal marker for transformation and expression) and merged signals *(merge*, merged YFP and RFP channels). Scale bars = 5 μm.

**Supplemental Fig. 6.**
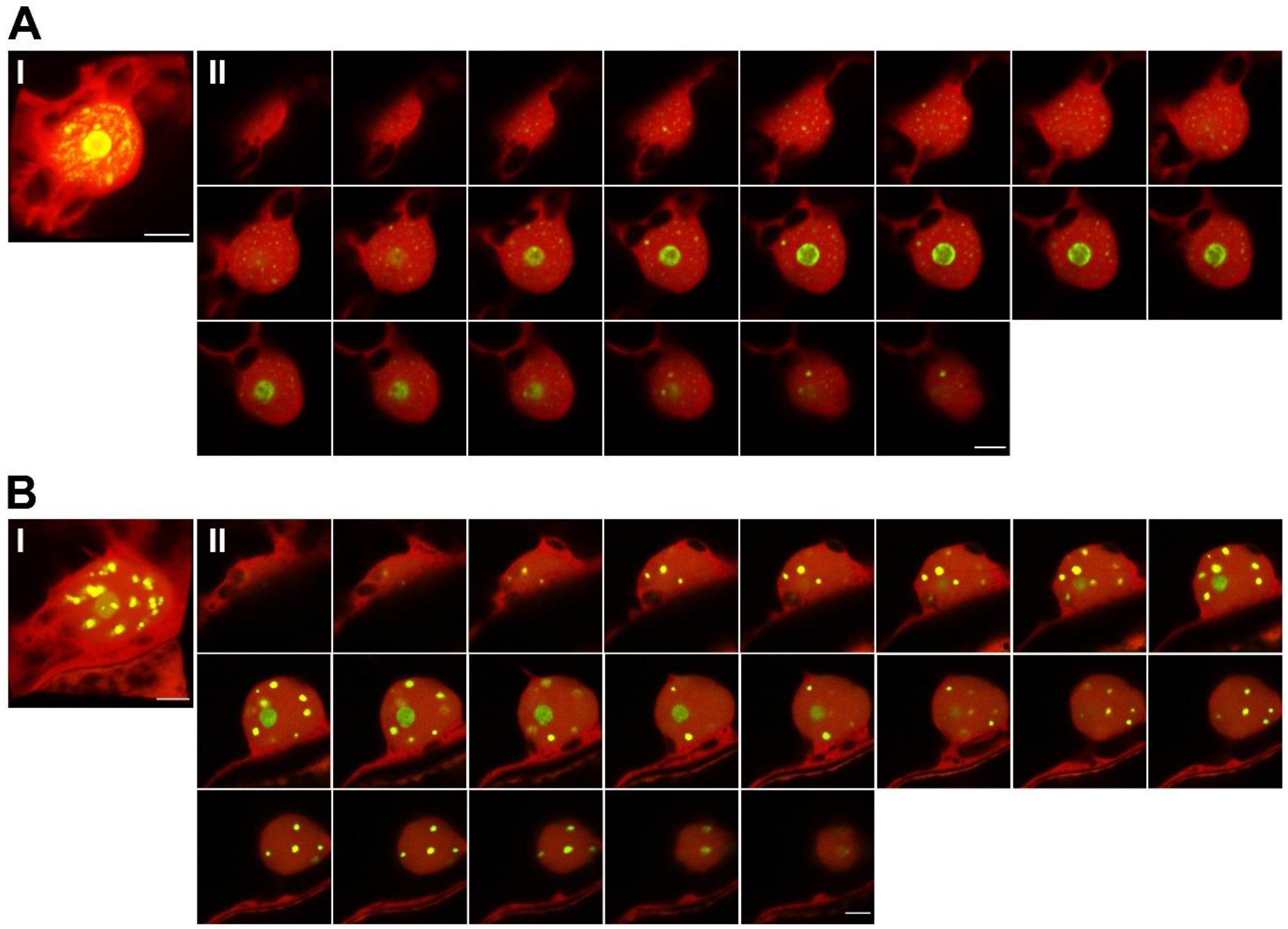
Maximum Intensity projections and Z-stacks of *N. benthamiana* epidermal cells nuclei presenting BiFC analyses. Maximum Intensity Projections (I) of entire Z-stack images (II) of nuclei of *N. benthamiana* leaf epidermal cells displaying BiFC interactions of TRB proteins. Shown here are merged images of YFP (interaction of the tested proteins) and mRFP (internal marker for transformation and expression) fluorescence detected by confocal microscopy. Scale bars = 5 μm.

A. Interaction of TRB1 and TRB2 proteins (nYFP-TRB1 + cYFP-TRB2 pBiFCt-2in1-NN construct). Optical sections 0,48 μm each.
B. Interaction of TRB1 and TRB4 proteins (nYFP TRB1 + cYFP TRB4 pBiFCt-2in1-NN construct). Optical sections 0,55 μm each.

**Supplemental Fig. 7.**
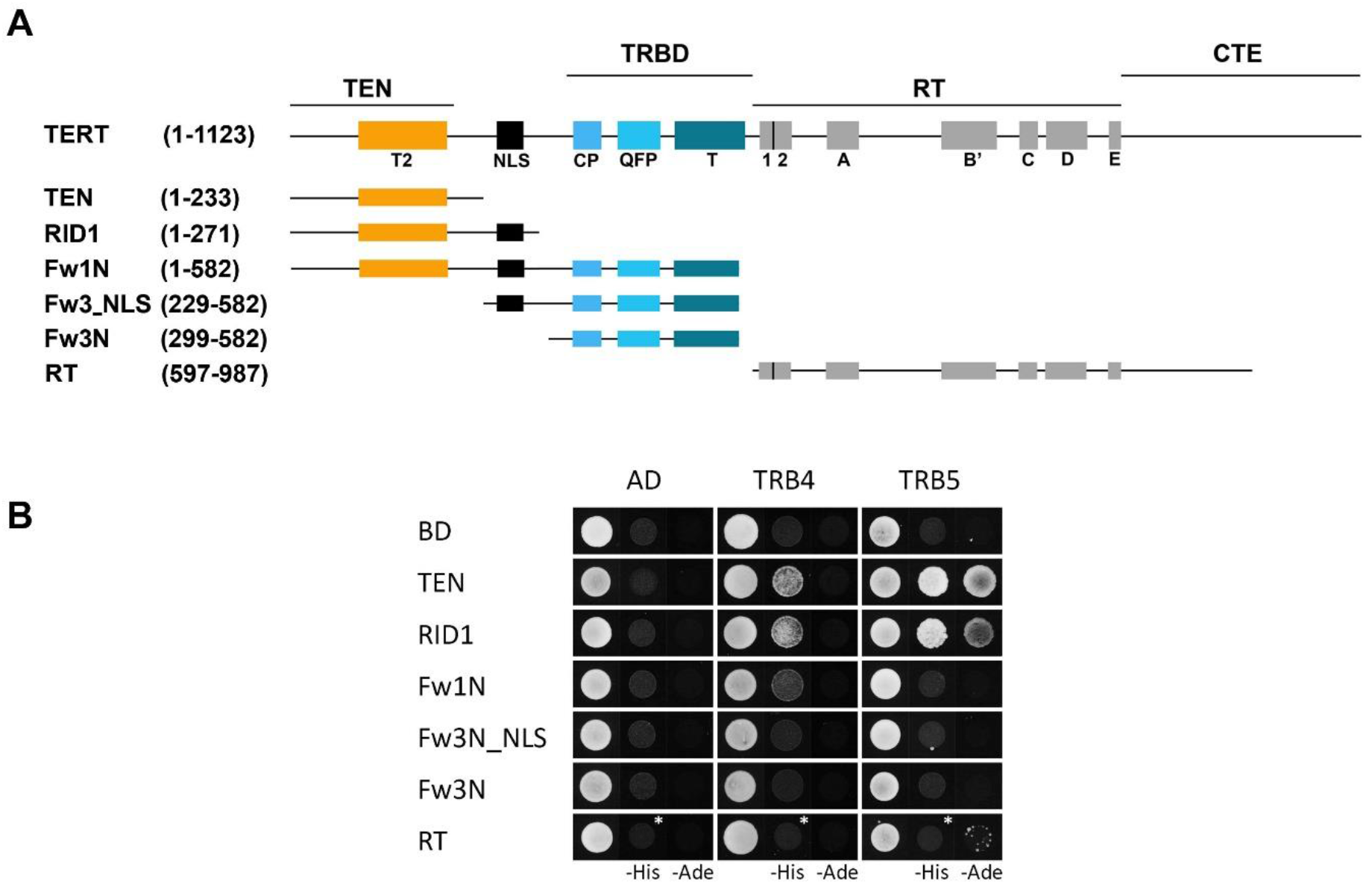
Interactions of TRB4-5 with TERT domains. A) Schematic depiction of the plant catalytic subunit of telomerase (TERT) showing functional motifs. The regions of structural domains TEN (telomerase essential N-terminal domain), TRBD (Telomerase RNA-binding domain), RT (reverse transcriptase domain) and CTE (C-terminal extension) are depicted above the conserved RT motifs (1, 2, A, B, C, D and E), telomerase-specific motifs (T2, CP, QFP and T) and a NLS (nucleus localization-like signal). All of the depicted TERT fragments were used for protein-protein interaction analysis (amino acid numbering is shown). B) TERT fragments from Majerská et al. 2017 were fused with the GAL4 DNA-binding domain (BD). TRB4 and TRB5 were fused with the GAL4 activation domain (AD). Both constructs were introduced into yeast strain PJ69-4a carrying reporter genes His3 and Ade2. Interactions were detected on histidine-deficient SD medium (-His), or under stringent adenine-deficient SD medium (-Ade) selection. Co-transformation with an empty vector (AD, BD) served as a negative control. *Asterisks* *, 3 mM 3-aminotriazol.

**Supplemental Fig. 8.**
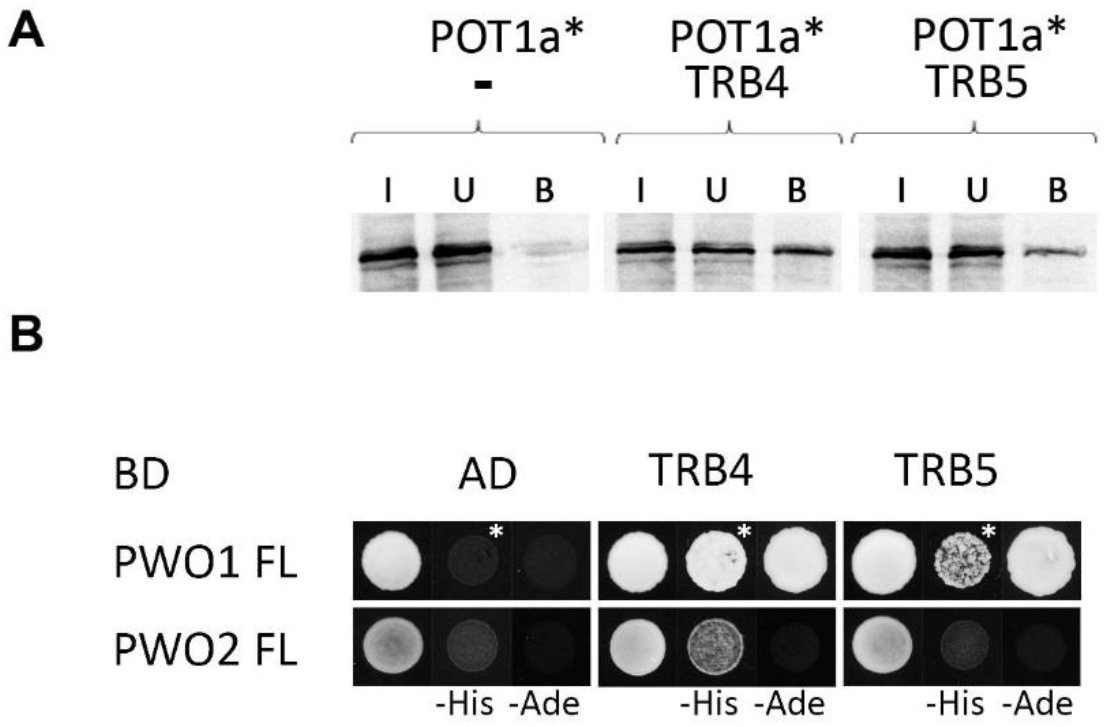
Protein-protein interactions of TRBs with various proteins. A) TNT expressed proteins, POT1a (^35^S-labelled*) and TRB4/TRB5 (myc-tag), were mixed and incubated with an anti-myc antibody and Protein G magnetic particles. In the control experiment, the POT1a proteins were incubated with an anti-myc antibody and Protein G magnetic particles in the absence of partner protein. Input (I), unbound (U), and bound (B) fractions were collected and run in SDS–10% PAGE gels. *Asterisks*\*, ^35^S labelling. B) Interactions of TRB4-5 were evaluated using the Y2H system. Interactions between TRB4-5 and full-length PWO1-2 were tested as in Fig. 7B. Interactions were detected on histidine-deficient SD medium (-His), or under stringent adenine-deficient SD medium (-Ade) selection. Co-transformation with an empty vector (AD, BD) served as a negative control. *Asterisks* *, 3 mM 3-aminotriazol.

Supplemental Table 1 List of the analyzed plant species for Fig. 1B and their accession numbers

Supplemental Table 2 List of the analyzed plant species for Fig. 2B and their accession numbers

Supplemental Table 3 List of the constructs tested by Bimolecular fluorescence complementation (BiFC) assay

TRB proteins were C-terminally (pBiFC 2in1-CC) and N-terminally (pBiFC 2in1-NN) fused to either nYFP or cYFP part. After transformation of *N. benthamiana* leaf epidermal cells with individual constructs, YFP fluorescence was observed: •, positive BiFC interaction, YFP signal observed in practically all transformed cells; ◦, weak YFP signal observed in few transformed cells only, most of the cells without YFP signal; x, no BiFC interaction, no YFP signal observed in any transformed cells.

Supplemental Table 4 List of primers and probes

